# Multi-target mode of action of Sulfodyne^®^, a stabilized Sulforaphane, against pathogenic effects of SARS-CoV-2 infection

**DOI:** 10.1101/2023.12.18.572126

**Authors:** Paul-Henri Romeo, Laurine Conquet, Sébastien Messiaen, Quentin Pascal, Stéphanie G. Moreno, Anne Bravard, Jacqueline Bernardino-Sgherri, Nathalie Dereuddre-Bosquet, Xavier Montagutelli, Roger Le Grand, Vanessa Petit, Federica Ferri

## Abstract

The coronavirus disease 2019 (COVID-19) due to the severe acute respiratory syndrome coronavirus 2 (SARS-CoV-2) has shown that, except vaccination, few therapeutics options for its treatment or prevention are available. Among the pathways that can be targeted for COVID-19 treatment, the Keap1/Nrf2 pathway seems of high interest as it regulates redox homeostasis and inflammation that are altered during SARS-CoV-2 infection. Here, we use three potent activators of the Keap1/Nrf2 pathway and showed that Sulfodyne^®^, a stabilized natural Sulforaphane preparation with optimal bioavailability, had the highest antiviral activity in pulmonary or colonic epithelial cell lines even when added late after SARS-CoV-2 infection. This antiviral activity was not dependent on NRF2 activity but associated with action on ER stress and mTOR signaling that are activated during SARS-CoV-2 infection. Sulfodyne^®^ also decreased the inflammatory response of epithelial cell lines infected by SARS-CoV-2 independently of SARS-CoV-2 replication and reduced the activation of human monocytes that are recruited after infection of epithelial cells by SARS-CoV-2. Administration of Sulfodyne^®^ had little effects on SARS-CoV-2 replication in mice and hamsters infected with SARS-CoV-2 but significantly reduced weight loss and disease severity. Altogether, these results pinpoint the natural compound Sulfodyne^®^ as a potent therapeutic agent of COVID-19 symptomatology.

**Author Summary:** Accumulating evidence shows that oxidative stress coupled with the systemic inflammation contribute to COVID-19 pathogenesis. As the Keap1/Nrf2 pathway is the major regulator of redox homeostasis and promotes resolution of inflammation and as lung biopsies from COVID-19 patients showed a decreased NRF2 target gene signature, pharmacological agents that are known to activate NRF2 are good candidates for COVID-19 treatment. We show herein that Sulfodyne^®^, an NRF2 activator that consists in a stabilized Sulforaphane preparation with optimal bioavailability, impairs SARS-CoV-2 replication in colonic or pulmonary epithelial cells. We show that this antiviral activity of Sulfodyne^®^ is not dependent of NRF2 activation, characterize the pathways associated with the Sulfodyne^®^ antiviral activity and show that Sulfodyne^®^ displays multiple actions that result in a decrease of the inflammation associated with SARS-CoV-2 infection. Finally, we show that Sulfodyne^®^ decreases the pathogenesis of mice or hamster infected with SARS-CoV-2. Overall, this study provides mechanistic explanations of the action of Sulfodyne^®^ during SARS-CoV-2 infection and suggests that Sulfodyne^®^ is a potential therapeutic agent of COVID-19 pathogenesis.

## Introduction

The most aggressive form of Acute Respiratory Distress Syndrome (ARDS) caused by SARS-CoV-2 is characterized by a cytokine storm and a leukopenia[1]. Whereas vaccination is the most effective prophylactic treatment against SARS-CoV-2 infection, other therapeutic approaches are still needed for patients that develop pathological effects after SARS-CoV-2 infection. Monoclonal antibodies, convalescent plasma, immunomodulators and antiviral drugs[2] are administered to patients with moderate/severe COVID-19 but most of them cannot be used at large-scale due to issues with expensive costs, route of administration and/or concerns about side effects. Thus, oral therapeutics with easily accessible drugs that decreased the symptomatology associated with SARS-CoV-2 infection are urgently required. The Keap1/Nrf2 pathway is the major redox-responsive pathway that protects cells against oxidative stress and damage and also represses pro-inflammatory cytokine genes[3–5]. In addition, NRF2 has antiviral properties that can be separated from its anti-inflammatory/cytoprotective properties[6]. As for many viruses, SARS-CoV-2 infection and replication are associated with an oxidative stress that predicts disease severity in infected patients. Both clinical and experimental evidence indicate that the Keap1/Nrf2 pathway is down-regulated by SARS-CoV-2. In accordance, analyses of lung biopsies from COVID-19 patients show that genes regulated by NRF2 have a decreased mRNA levels[7] and expression of NRF2 protein is decreased in children infected with SARS-CoV-2[8]. Furthermore, the NSP14 protein of SARS-CoV-2 impairs NRF2/HMOX1 activation[9], the SARS-CoV-2 ORF3a positively regulates ferroptosis by degradation of NRF2[10] and SARS-CoV-2 directly inhibits NRF2-mediated antioxidant response[11]. Interestingly, SARS-CoV-2 replication does not seem to be dependent on the Keap1/Nrf2 pathway but the pathogenic consequences of SARS-CoV-2 infection depend on the Keap1/Nrf2 pathway[11]. These results indicate that activating the Keap1/Nrf2 pathway may be a therapeutic approach during SARS-CoV-2 infection and thus could at least complement an antiviral therapy.

There are numerous NRF2 activating drugs, used in clinical trials, that are good candidates for treating the pathogenic consequences of SARS-CoV-2 infection. PB125^®^, a phytochemical NRF2-activating ingredient, inhibits SARS-CoV-2 entry by downregulating ACE2 and TMPRSS2, two SARS-CoV-2 entry factors and downregulates several inflammatory cytokines that are known markers of severe COVID-19, in endotoxin-stimulated cells[12]. Dimethyl fumarate (DMF) and 4-octyl-itaconate (4-OI) inhibit viral replication in epithelial cells and decrease the expression of associated inflammatory genes when used in prophylactic treatment *in vitro*[7]. Bardoxolone methyl (CDDO-Me) possesses an anti-inflammatory activity against viruses like DENV or Zika virus[13]. Finally, Sulforaphane (SFN), an orally accessible and well-tolerated dietary supplement found in high concentration in broccoli and other cruciferous vegetables, inhibits SARS-CoV-2 replication in cell lines and displays anti-inflammatory action during SARS-CoV-2 infection[14]. However, the antiviral activity of SFN in epithelial cell lines is conserved in NRF2-knockdown cells[14] and its action may be related to its reversible inhibition of 3-chymotrypsin-like protease of SARS-CoV-2[15], its inhibition of the NF-kB pathway[16] and/or its inhibitory effect on NRLP3 inflammasome activation[17].

We report here that the natural compound Sulfodyne^®^, a stabilized SFN preparation that provides optimal bioavailability[18], has a more efficient antiviral activity against SARS-CoV-2 than DMF or CDDO. We characterized the pathways associated with this antiviral activity, studied the relationship between SARS-CoV-2 replication and the anti-inflammatory action of Sulfodyne^®^, characterized the by-stander effects of Sulfodyne^®^ on human monocytes and showed that, *in vivo*, Sulfodyne^®^ decreased the pathological consequences of SARS-CoV-2 infection despite little action on viral replication.

## Results

### Increased susceptibility of *Nrf2* KO mice to SARS-CoV-2 infection

Respiratory viral infections, including SARS-CoV-2 infection, result in inflammation and oxidative injury. To characterize the role of NRF2, the master regulator of the antioxidant response, during SARS-CoV-2 infection *in vivo*, we used mice genetically deficient in NRF2 (*Nrf2* KO). *Nrf2* KO and C57BL/6 wild-type (WT) mice were infected with Beta SARS-CoV-2 variant (Fig. S1A) and clinical development of disease was monitored for 14 days post-infection (dpi). Quantification of viral load in the lung showed higher levels in *Nrf2* KO than in WT mice at 6dpi (Fig. S1B), and *Nrf2* KO mice displayed an increased loss of body weight and a longer time to return to their pre-infection body weight (Fig. S1C). Furthermore, whereas WT mice did not show any clinical symptom, *Nrf2* KO mice exhibited signs of pathological infection (measured by ruffled fur, hunched posture, reduced locomotion and difficult breathing) with maximal score at 3dpi, indicating an increased susceptibility of *Nrf2* KO mice to SARS-CoV-2-induced disease (Fig. S1D). Finally, histopathological analysis of lungs at 6dpi showed an increase in pulmonary lesions and a more severe bronchial inflammation in *Nrf2* KO mice compared to WT (Fig. S1E).

Altogether, these results indicate an increased pathological response of *Nrf2* KO mice to SARS-CoV-2 infection and suggest that NRF2 contributes to the antiviral response against SARS-CoV-2 and/or to the pathological effects associated with SARS-CoV-2 infection.

### Sulfodyne^®^ exhibits strong antiviral activity in epithelial cell lines

Results from *Nrf2* KO mice prompted us to investigate the effect of pharmacological activation of NRF2 on SARS-CoV-2 infection. Using Calu-3 cells, a lung human epithelial cell line, highly permissive for SARS-CoV-2 infection, we studied the effect of three NRF2 agonists: the clinically approved Dimethyl Fumarate (DMF), the synthetic triterpenoid CDDO-Imidazolide (CDDO) and the natural compound Sulfodyne^®^, a stabilized Sulforaphane (SFN) preparation that provides optimal bioavailability. First, we showed that Sulfodyne^®^ treatment induced NRF2 nuclear translocation in Calu-3 cells when used at a concentration equivalent to 14uM of SFN (Fig. S2A) without affecting cell viability (Fig. S2B).

To evaluate the antiviral activity of these NRF2 agonists, Calu-3 cells were infected with the initial pandemic SARS-CoV-2 Wuhan strain (thereafter referred as SARS-CoV-2) at MOI 0.5, for 12h before drugs addition (Fig. 1A, left panel). All drugs activated NRF2, as illustrated 36 hours post infection (hpi) by the increased mRNA level of HMOX1, a known NRF2 target gene (Fig. 1A, middle panel). Quantification of intracellular genomic viral RNA expression showed that Sulfodyne^®^ was a potent suppressor of SARS-CoV-2 infection of Calu-3 cells, whereas DMF and CDDO displayed lower antiviral activity (Fig. 1A, right panel). In addition, Sulfodyne^®^ but neither DMF nor CDDO decreased SARS-CoV-2 infection in the human colonic epithelial Caco2 cells whereas all drugs activated the Keap1/Nrf2 pathway (Fig. 1B). We thus focused our studies on the effect of Sulfodyne^®^ on SARS-CoV-2 infection.

**Fig.1.**
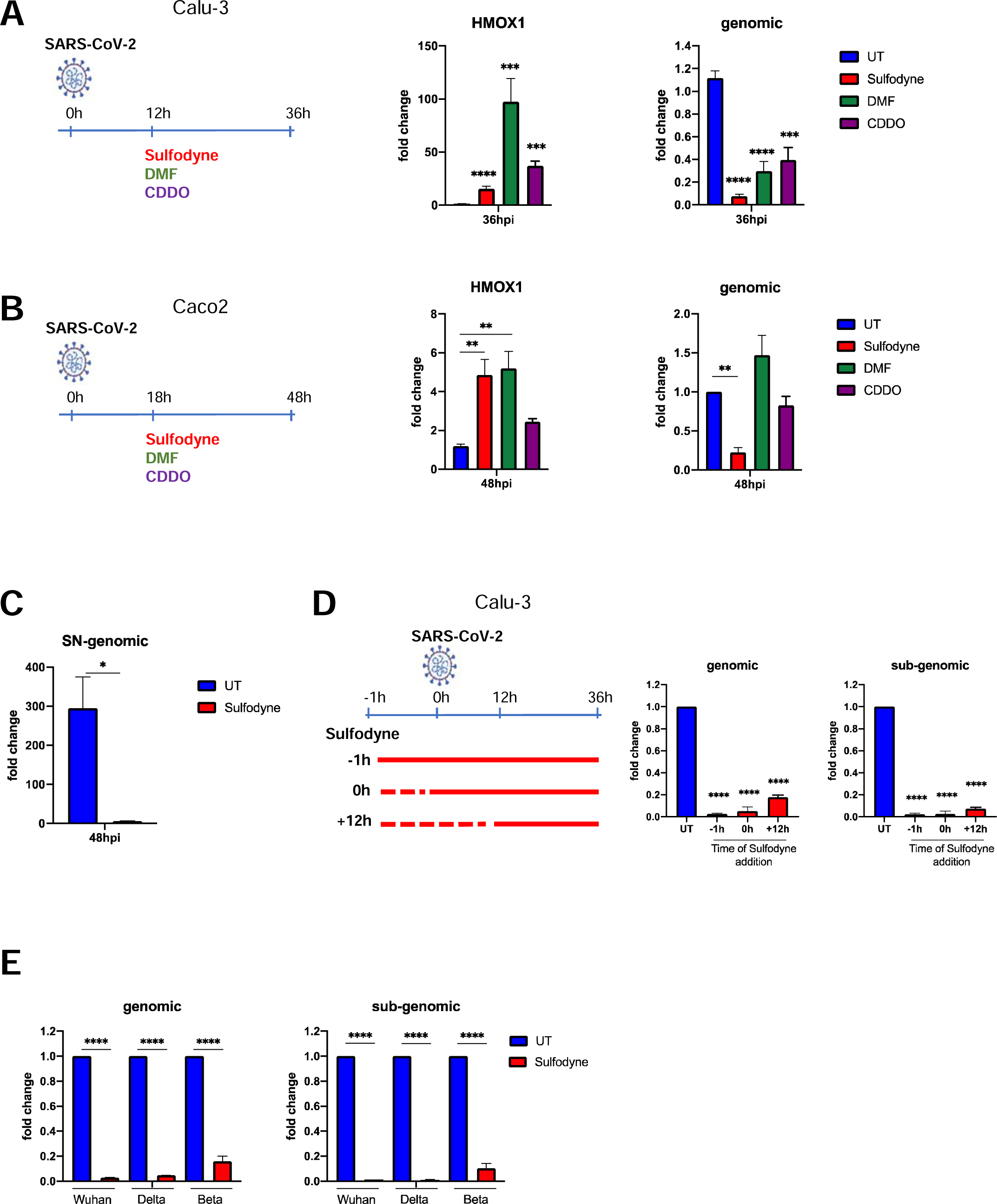
Sulfodyne^®^ efficiently inhibits SARS-CoV-2 replication in epithelial cells A) Calu-3 and B) Caco2 cells were infected with SARS-CoV-2 (Wuhan) at MOI of 0.5 and were untreated (UT) or treated with indicated NRF2 activators at 12 or 18 hours post-infection (hpi). Viral genomic RNA (primers targeting the RdRp gene in ORF1) and cellular *HMOX1* (a known NRF2 target gene) mRNA were quantified 36hpi (Calu-3) or 48hpi (Caco2) by RT-qPCR. Data are expressed as fold change over the UT. Mean ± sem. Calu-3: n=11 for UT, Sulfodyne^®^ and DMF; n=2 for CDDO. Caco2: n=6 for UT, Sulfodyne^®^ and DMF; n=2 for CDDO. One-way ANOVA with Dunnett’s multiple comparisons test vs UT. B) Expression of viral genomic RNA at 48hpi in supernatant (SN) from Calu-3 cells infected with SARS-CoV-2 at MOI of 0.5 and untreated (UT) or treated with Sulfodyne^®^ at 12hpi. Data are expressed as fold change over the inoculum. Mean ± sem. n=3. Unpaired t-test. C) Time-of-Sulfodyne^®^ treatment-assay. Calu-3 cells were infected with SARS-CoV-2 at MOI of 0.5 and treated with Sulfodyne^®^ at indicated times before, at the time or after infection. Viral genomic and sub-genomic (primers spanning the Leader sequence and the E-gene) RNA was quantified at 36hpi and expressed as fold change over the UT. Mean ± sem; n=2. One-way ANOVA with Dunnett’s multiple comparisons test vs UT at 36hpi. D) Calu-3 were infected with Wuhan and variant of concern Delta and Beta SARS-CoV-2 at MOI of 0.5, treated with Sulfodyne^®^ at 12hpi and collected at 36hpi for genomic and sub-genomic RNA quantification. Data are shown relative to expression of UT at 36hpi. Mean ± sem; n=2. One-way ANOVA with Sidak’s multiple comparisons test vs the corresponding UT at 36hpi. * p ≤ 0.05; ** p≤ 0.01; *** p ≤ 0.001; **** p≤ 0.0001.

The levels of genomic viral RNA load in culture supernatants of Calu-3 cells showed that the decreased intracellular viral RNA after Sulfodyne^®^ treatment was associated with a decreased SARS-CoV-2 spread (Fig. 1C). If added prior to, at the time of, or after SARS-CoV-2 infection, Sulfodyne^®^ activated the Keap1/Nrf2 pathway (Fig. S2C) and was highly effective (>95% of inhibition) in reducing genomic and sub-genomic SARS-CoV-2 expression (Fig. 1D). We finally studied the antiviral activity of Sulfodyne^®^ against the Delta and Beta SARS-CoV-2 variants. Sulfodyne^®^ treatment decreased viral genomic and sub-genomic expression in Calu-3 cells infected with Delta and Beta strains with comparable efficacy to that reported to reference strain Wuhan (Fig. 1E), indicating that Sulfodyne^®^ may inhibit replication of SARS-CoV-2 variants.

Altogether, these results showed a strong antiviral activity of Sulfodyne^®^ against SARS-CoV-2 infections in pulmonary or colonic epithelial cell lines.

### Sulfodyne^®^ inhibits SARS-CoV-2 replication

Whereas NRF2 deficiency increases ACE2 expression[19], activation of NRF2 decreases ACE2 expression[12, 19] and increases thioredoxin reductase (TRXR) which could reduce the interaction between SARS-CoV-2 Spike protein and ACE2 and thus decreases SARS-CoV-2 entry[20].

To investigate if the antiviral activity of Sulfodyne^®^ was due to an impaired SARS-CoV-2 entry process to host cells, we studied the effect of Sulfodyne^®^ on cell-surface expression of ACE2, the main SARS-CoV-2 receptor in Calu-3 cells. No significant change in the percentage of ACE2-expressing Calu-3 cells nor in the ACE2 expression level at the surface of ACE2-positive cells was observed after Sulfodyne^®^ treatment (Fig. 2A). To further characterize the Sulfodyne^®^ mode of action, Calu-3 cells were infected with SARS-CoV-2 for 12h before Sulfodyne^®^ treatment, in presence or absence of an anti-SARS-CoV-2 Spike antibody that blocks viral entry (Fig. S3A) and thus prevents reinfection. Quantification of intracellular viral RNA at 36hpi showed a similar decrease in both genomic and sub-genomic RNA in the presence or absence of anti-Spike antibody in Sulfodyne^®^-treated compared to untreated cells (Fig. 2B). These results indicate that Sulfodyne^®^ did not modify SARS-CoV-2 entry in Calu-3 cells.

**Fig.2.**
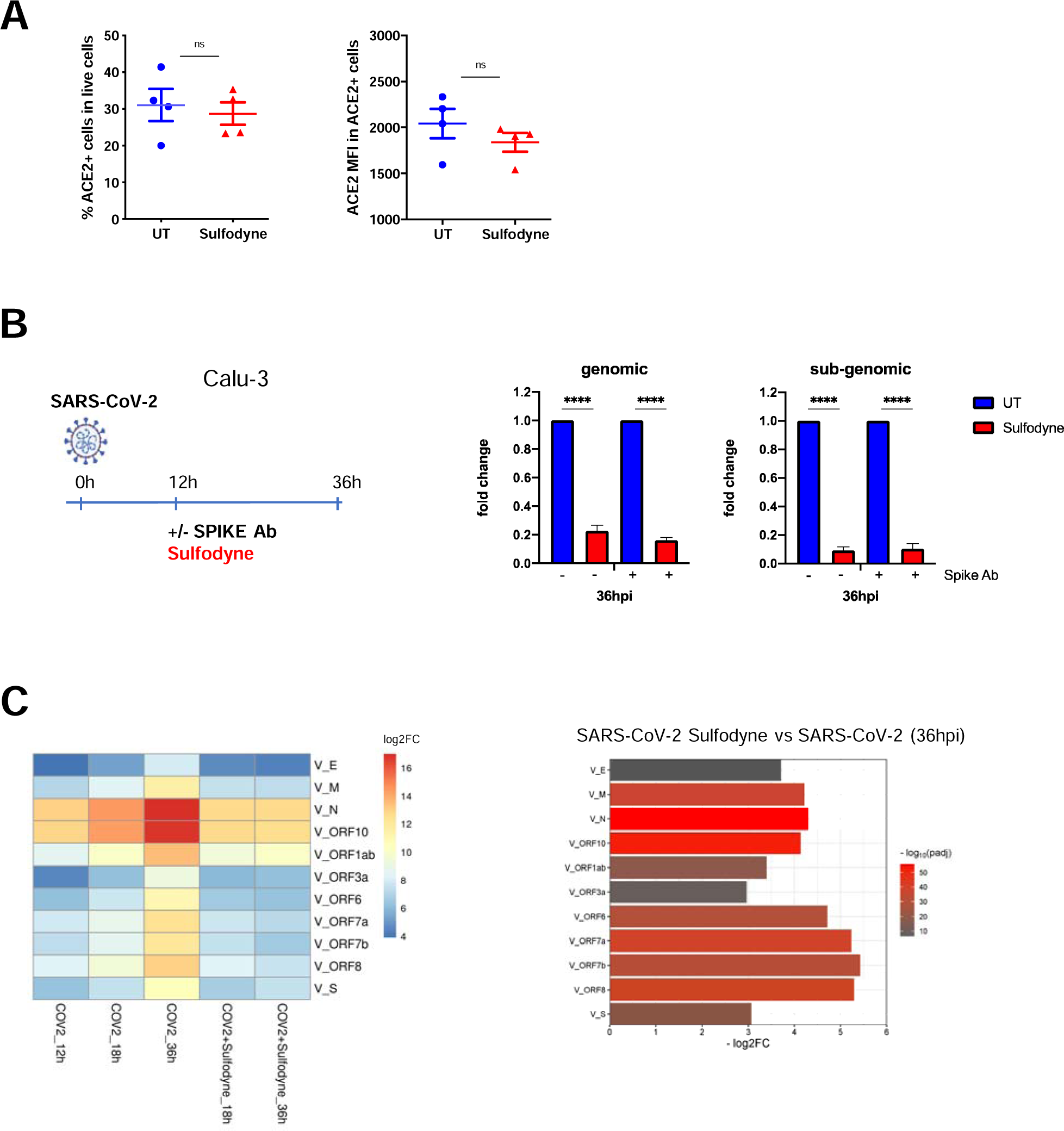
Sulfodyne^®^ blocks SARS-CoV-2 replication at SARS-CoV-2 post-entry stage A) Calu-3 cells were untreated (UT) or treated with Sulfodyne^®^ for 48h and ACE2 expression at the membrane of the Calu-3 cells (Left) and ACE2 mean fluorescence intensity (MFI) in ACE2 positive cells (right) were studied by flow cytometry. Mean ± sem; n=4. Two-tailed Mann-Whitney U test, ns: not significant B) Calu-3 cells were infected with SARS-CoV-2 for 12 hours. SARS-CoV-2 was then washed, Sulfodyne^®^ was added in presence of the anti-Spike neutralizing antibody (Spike Ab) and genomic and sub-genomic viral RNA were quantified at 36hpi. Data are presented as fold decrease relative to the corresponding UT sample. Mean ± sem; n=3. One-way ANOVA with Sidak’s multiple comparisons test vs corresponding UT at 36hpi. **** p≤ 0.0001. C) (Left) Heatmap showing log2FC of indicated RNA-seq samples compared to mock Calu-3 cells for each individual SARS-CoV-2 transcript. Sulfodyne^®^ was added at 12hpi. n=4. (Right) Fold decrease of viral transcripts at 36hpi in Calu-3 cells infected SARS-CoV-2 and treated with Sulfodyne^®^ compared to SARS-CoV-2 infected cells. Colours represent p-adj from DEseq2 analysis.

To characterize the kinetics of inhibition of SARS-CoV-2 replication by Sulfodyne^®^, RNA-seq experiment was performed over a time course of 12h, 18h and 36h in Calu-3 cells infected with SARS-CoV-2 and treated with Sulfodyne^®^ at 12hpi. The number of reads mapping to SARS-CoV-2 genes increased over time, with a log2 fold-change (FC) of 4-13 at 12hpi, a log2FC of 5-14 at 18hpi and a log2FC of 8-17 at 36hpi, indicating an active replication (Fig. 2C). Sulfodyne^®^ treatment at 12hpi completely blocked transcription of all viral transcripts over time (Fig. 2C left panel), with a log2 fold decrease of 3-5 at 36hpi compared to SARS-CoV-2 infected Calu-3 cells (Fig. 2C right panel). These results indicate that Sulfodyne^®^ completely inhibited SARS-CoV-2 replication in Calu-3 cells.

### Sulfodyne^®^ regulates host metabolic pathways during SARS-CoV-2 infection

Few transcriptional changes of endogenous genes were observed in Calu-3 cells 12h and 18h after SARS-CoV-2 infection (respectively 15 and 36 differentially expressed genes, p-adj<0.05), whereas 500 differentially expressed genes (DEG, p-adj<0.05) were identified at 36hpi. Sulfodyne^®^ reversed expression of genes up-regulated by SARS-CoV-2 infection at 36dpi (Fig. S4A), that are mainly genes associated to interferon (IFN) and antiviral defence pathways (Fig. S4B, upper panel). Genes down-regulated by SARS-CoV-2 infection were mainly associated with the mitochondrial electron transport chain/oxidative phosphorylation (Fig. S4B, lower panel). Sulfodyne^®^ treatment reversed expression of about 35% of genes that were down-regulated by SARS-CoV-2 infection at 36dpi (Fig. S4A), suggesting a partial effect of Sulfodyne^®^ on oxidative phosphorylation.

The expression of genes linked to the NRF2 pathway was not significantly modified during the first 36 hours of Calu-3 cells infection by SARS-CoV-2 (Fig. 3A). By contrast, Sulfodyne^®^ treatment during infection led to a significant induction of ARE-containing cytoprotective genes, coding for key components in antioxidant systems (Glutathione- and thioredoxin-based systems; heme and iron metabolism), drug detoxification (NQO1; UDP-glucuronosyltransferase UGT), chaperones involved in protein folding (HSP90AA1, HSPA1A) and components of proteasome (PSMD3, PSMD4) (Fig. 3A).

**Fig.3.**
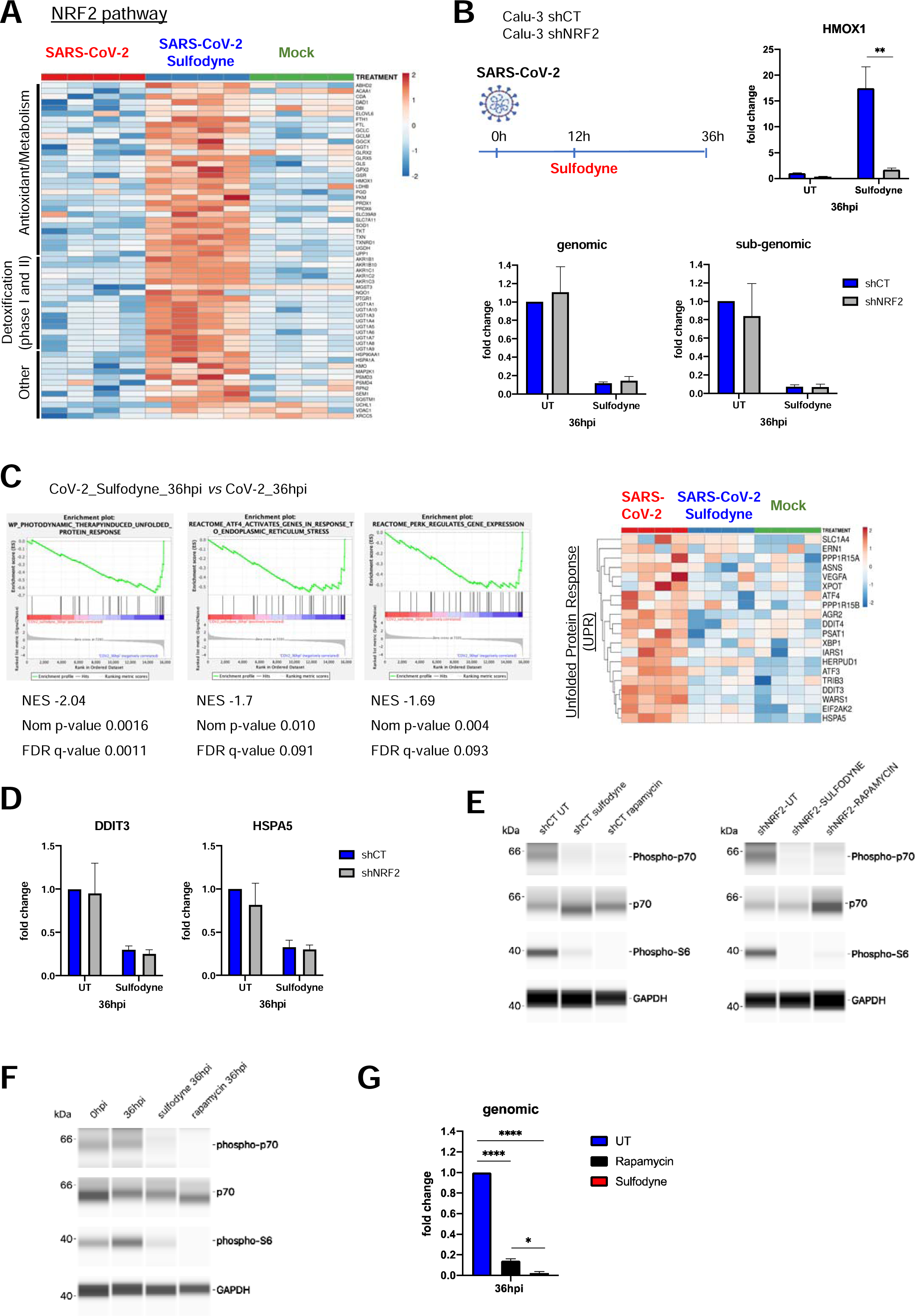
Characterization of the pathways regulated by Sulfodyne^®^ in Calu-3 cells infected with SARS-CoV-2 A) Heatmap of normalized expression levels of DEG (SARS-CoV-2 +Sulfodyne^®^ vs SARS-CoV-2 at 36hpi) related to the Keap1/NRF2 pathway in Calu-3 cells. Normalized counts are log-transformed, centered and scaled by row. The columns display the data for each of the 4 replicates. B) shCT and shNRF2 Calu-3 cells were infected with SARS-CoV-2 and treated with Sulfodyne^®^ at 12hpi. *HMOX1* and viral RNA expression were quantified at 36hpi and shown relative to the UT shCT. Mean ± sem; n=3. Two-way ANOVA with Sidak’s multiple comparisons test vs corresponding shCT. C) (Left) Gene set enrichment analysis (GSEA) showing ER stress/UPR-related pathways that are negatively correlated in SARS-CoV-2+Sulfodyne^®^ (red) *vs* SARS-CoV-2 (blue) at 36hpi. NES, normalized enrichment score. Nom p-value= nominal p-value. FDR, False Discovery Rate. (Right) Heatmap of normalized expression levels of DEG related to the UPR. D) *DDIT3* and *HSPA5* mRNA levels at 36hpi in shCT and shNRF2 Calu-3 infected and treated with Sulfodyne^®^ at 12hpi. Data are expressed as fold change over UT shCT. Mean ± sem; n=3 E) WES Simple assay for mTOR substrate proteins phospho-p70 S6 kinase (Thr389; phospho-p70), p70 S6 kinase (p70), phospho-S6 ribosomal protein (Ser235/236; phospho-S6) and GAPDH in total extracts from shCT and shNRF2 Calu-3 cells treated with Sulfodyne^®^ or Rapamycin for 6h. F) WES Simple assay for mTOR substrate proteins and GAPDH in total extracts from Calu-3 cells non infected (0hpi), infected with SARS-CoV-2 (36hpi) or infected for 12h and then treated with Sulfodyne^®^ (Sulfodyne^®^ 36hpi) or Rapamycin (Rapamycin 36hpi). G) SARS-CoV-2 genomic RNA levels in Calu-3 cells infected and treated as in (f), relative to UT at 36hpi. Mean ± sem; n=2. One-way ANOVA with Tukey’s multiple comparisons test. * p ≤ 0.05; ** p≤ 0.01; *** p ≤ 0.001; **** p≤ 0.0001.

To characterize the role of NRF2 in the antiviral action of Sulfodyne^®^, NRF2 expression was down-regulated in Calu-3 cells using shRNA (Fig. S4C). As expected, NRF2 knock-down impaired the increased HMOX1 mRNA levels observed after Sulfodyne^®^ treatment (Fig. 3B, upper right panel). In accordance with the transcriptomic data, no significant effect on SARS-CoV-2 replication in Calu-3 cells was observed following NRF2 knock-down (Fig. 3B, lower panels) but, surprisingly, NRF2 knock-down did not modify the decrease in genomic and sub-genomic RNA expression observed after Sulfodyne^®^ treatment of SARS-CoV-2 infected Calu-3 cells (Fig. 3B, lower panels). Similar results were obtained when NRF2 was knocked-down in Caco2 cells (Fig. S4D and E). These results indicate that NRF2 activation by Sulfodyne^®^ treatment is not required for its antiviral action and prompted us to characterize pathways regulated by Sulfodyne^®^ and involved in its antiviral action.

Transcriptomic analyses at 36hpi revealed a significant enrichment of pathways related to endoplasmic reticulum (ER) stress and the unfolded protein response (UPR) among genes that are up-regulated during SARS-CoV-2 infection and down-regulated after Sulfodyne^®^ treatment (Fig. 3C). mRNA levels of UPR genes, including the UPR initiation marker HSPA5 (also kown as BIP/GPR78) and critical transcription factors such as XBP1, ATF4 and DDIT3, were increased after SARS-CoV-2 infection and these increases were reverted after Sulfodyne^®^ treatment at 12hpi (Fig. 3C, right panel). This Sulfodyne^®^-dependent inhibition of HSPA5 and DDIT3 increase during SARS-CoV-2 infection was also observed in NRF2 knock-down Calu-3 cells (Fig. 3D), indicating a NRF2-independent action of Sulfodyne^®^ on the ER stress.

It has been suggested that SARS-CoV-2 activates the mTOR (mammalian Target Of Rapamycin) pathway during infection[21]. We therefore studied if Sulfodyne^®^ could act on the mTOR pathway to block SARS-CoV-2 replication. As a hallmark of mTOR activation, we investigated phosphorylation of p70S6K (Thr389) and pS6 (Ser235/236), two ribosomal proteins essential for protein synthesis. Sulfodyne^®^ treatment decreased phosphorylation of these proteins in an NRF2-independent manner and this decrease was identical to that obtained after action of Rapamycin, the main inhibitor of mTOR (Fig. 3E). Sulfodyne^®^ treatment of Calu-3 or Caco2 cells after SARS-CoV-2 infection led to a decreased phosphorylation of p70S6k and pS6 similar to the one observed after treatment with Rapamycin (Fig. 3F and Fig. S4F), indicating that Sulfodyne^®^ inhibits the mTOR pathway during SARS-CoV-2 infection. Rapamycin treatment of Calu-3 cells decreased SARS-CoV-2 replication but less than Sulfodyne^®^ treatment (Fig. 3G). Altogether, these results indicate that, although the antiviral action of Sulfodyne^®^ in Calu-3 or Caco2 cells is not dependent of NRF2 expression, Sulfodyne^®^ treatment can repress SARS-CoV-2 replication by its action on several protective cellular metabolic pathways activated during SARS-CoV-2 infection, such as ER stress and mTOR signalling, and may restore redox homeostasis through activation of NRF2 target genes.

### Anti-inflammatory action of Sulfodyne^®^ is independent of SARS-CoV-2 replication

In accordance with previous results[22–24], we found that Calu-3 cells activate delayed IFN and antiviral defence responses relative to SARS-CoV-2 replication (Fig. 4A and Fig. 4B, upper panel), which might also contribute to pathogenic inflammatory response. Sulfodyne^®^ treatment 12h post-infection dampened type I IFN responses (Fig. 4B, upper panel) and mRNA levels of several inflammatory genes (Fig. 4B, lower panel), including genes encoding for chemo-attractants for monocytes and neutrophils. As for its action on viral infection, the anti-inflammatory activity of Sulfodyne^®^ was independent of NRF2 expression (Fig. 4C).

**Fig.4.**
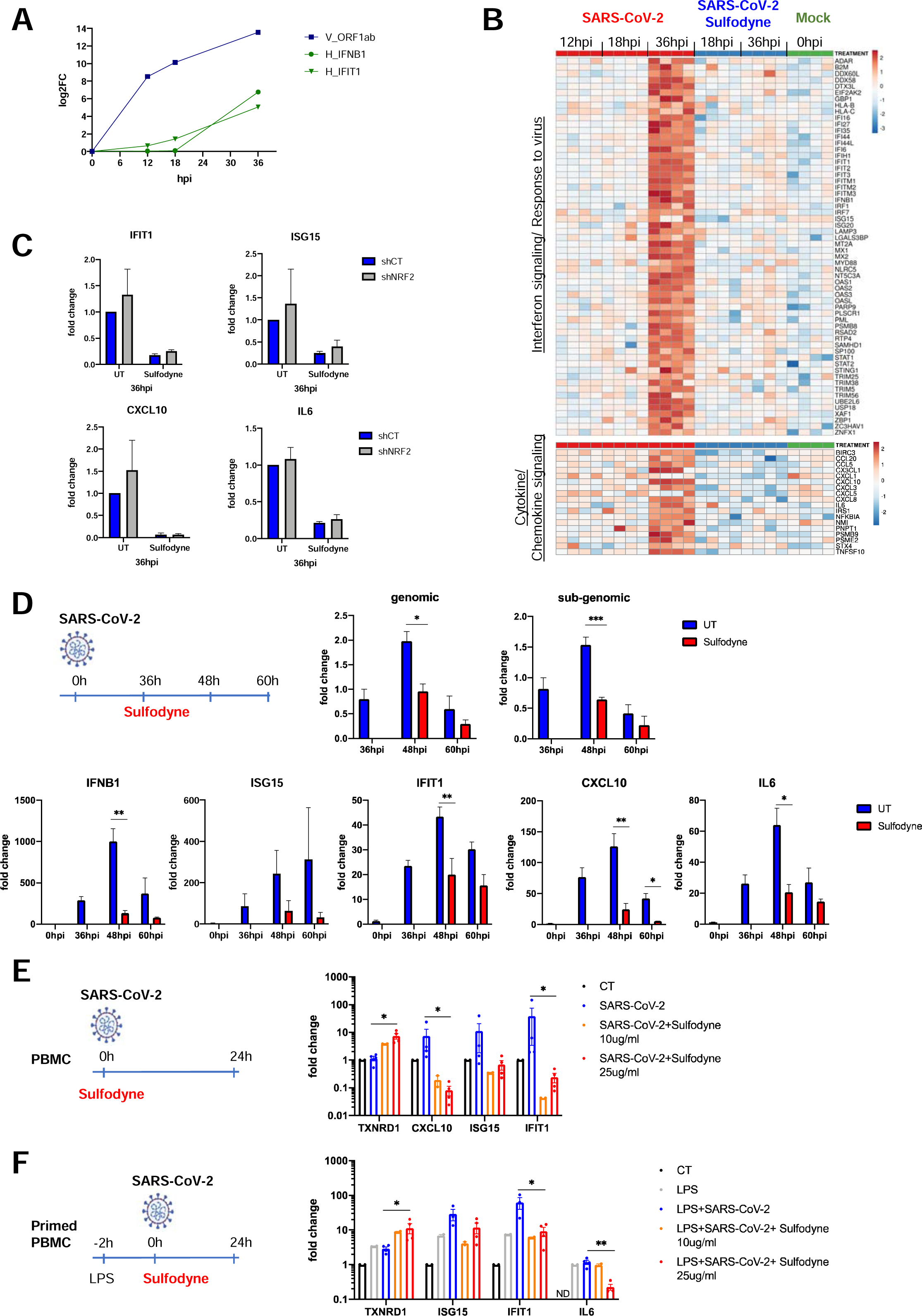
Anti-inflammatory actions of Sulfodyne^®^ A) Kinetics of viral replication (ORF1ab SARS-CoV-2 gene) and of the mRNA levels of the cellular genes *IFNB1* and *IFIT1* (IFN stimulated gene) in Calu-3 cells infected with SARS-CoV-2. Data are the mean of 4 replicates from RNA-seq experiment and presented as Log2FC over the mock Calu-3 cells. B) Heatmap of normalized expression levels of DEG (SARS-CoV-2 +Sulfodyne^®^ vs SARS-CoV-2 at 36hpi) related to Interferon signaling and Response to virus (upper panel) and Cytokine and Chemokine signaling pathways (lower panel) for all RNA-seq samples in Calu-3 cells. C) mRNA levels of the indicated genes in shCT and shNRF2 Calu-3 cells infected with SARS-CoV-2 and treated with Sulfodyne^®^ at 12hpi. Data are presented relative to UT shCT at 36hpi. Mean ± sem, n=3. D) Kinetics of viral RNA expression (upper panel) and of the mRNA levels of the indicated interferon-stimulated and inflammatory genes (lower panel) in Calu-3 cells infected with SARS-CoV-2 and treated with Sulfodyne^®^ at 36hpi. Data are presented relative to the UT cells at 36hpi for viral RNA and relative to uninfected cells (0hpi) for host genes. Mean ± sem; n=3. One-way ANOVA with Sidak’s multiple comparisons test UT vs Sulfodyne^®^ at each time. E) Human PBMC from healthy donors were mock-infected (CT), infected with SARS-CoV-2 in presence or not of indicated concentration of Sulfodyne^®^ for 24h and analysed for mRNA levels of the indicated genes. Data are expressed as fold change over CT PBMC. Mean ± sem, n=2-4. Two-tailed Mann-Whitney U test (SARS-COV-2+ Sulfodyne^®^ 25ug/ml *vs* SARS-COV-2). F) Human PBMC were primed for 2h with LPS before being infected and treated as in (E) and analysed for mRNA levels of indicated genes. Data are expressed as fold change over PBMC. Mean ± sem, n=2-4. Two-tailed Mann-Whitney U test (LPS+SARS-COV-2+ Sulfodyne^®^ 25ug/ml *vs* LPS+SARS-COV-2). ND= not detected. * p ≤ 0.05; ** p≤ 0.01; *** p ≤ 0.001

As SARS-CoV-2 replication induced an inflammatory response in Calu-3 cells, the observed decreased mRNA levels of inflammatory genes after Sulfodyne^®^ treatment might be a consequence of decreased viral replication. To test this hypothesis, Calu-3 cells were treated with Sulfodyne^®^ at 36hpi, a time at which Calu-3 inflammatory response is high (Fig. 4B) and active viral replication is low (Fig. 4A and Fig. 4D, upper panels). Kinetics analysis showed that Sulfodyne^®^ decreased mRNA levels of IFNB1, IFN stimulated genes (ISG15, IFIT1), CXCL10 and IL6 (Fig. 4D, lower panels), suggesting that the anti-inflammatory activity of Sulfodyne^®^ is not only due to decreased SARS-CoV-2 replication.

Peripheral blood immune cells, recruited to the lung compartment, are major contributors to human inflammatory responses after SARS-CoV-2 infection. Infection of human peripheral blood mononuclear cells (PBMC) with SARS-CoV-2 is not productive as SARS-CoV-2 cannot replicate in PBMC, but exposure of PBMC to SARS-CoV-2 can induce an innate immune response[25]. We therefore studied the effects of Sulfodyne^®^ treatment on inflammatory response of PBMC exposed to SARS-CoV-2. PBMC from healthy donors were exposed *ex vivo* to SARS-CoV-2 and treated with different doses of Sulfodyne^®^ for 24h (Fig. 4E). Sulfodyne^®^ activated NRF2 in a dose-dependent manner, as measured by mRNA level of TXNRD1, a known NRF2 target gene and decreased mRNA levels of CXCL10 and interferon-stimulated genes IFIT1 and ISG15 which were increased after SARS-CoV-2 infection (Fig. 4E).

Furthermore, as SARS-CoV-2 infection of immune cells *in vivo* can occur in a pre-existing inflammatory environment, PBMC were primed with LPS 2h before viral infection and Sulfodyne^®^ treatment (Fig. 4F, left panel). Under this condition, Sulfodyne^®^ activated NRF2 (Fig. 4F, right panel). Exposure of PBMC to LPS alone increased IFIT1 and ISG15 mRNA levels that were further increased following SARS-CoV-2 infection. Sulfodyne^®^ treatment only reduced the viral-dependent increase of these mRNA levels. In contrast, IL6 mRNA level was not increased after SARS-CoV-2 infection and high concentration of Sulfodyne^®^ decreased IL6 mRNA level (Fig. 4F, right panel).

Altogether these results indicated that, in presence of Sulfodyne^®^, the decreased inflammatory response associated with SARS-CoV-2 infection in both epithelial cells and primary immune cells is not dependent on SARS-CoV-2 replication.

### Immunomodulatory impact of Sulfodyne^®^ on by-stander monocytes

High release of inflammatory mediators by infected lung epithelial cells may exacerbate monocytes infiltration and immune response, leading to severe COVID-19 pathological effect and the anti-inflammatory action of Sulfodyne^®^ in lung epithelial cells could be beneficial in preventing hyperinflammatory reactions that contribute to the lung damage.

We therefore assessed if SARS-CoV-2 infection of lung epithelial cells can promote an inflammatory response in human CD14^+^ monocytes and studied the effect of Sulfodyne^®^ treatment on this inflammatory response. Human CD14^+^ monocytes were purified from healthy donors and cultured for 2h and 8h in presence of supernatant from uninfected, SARS-CoV-2-infected, and SARS-CoV-2-infected and Sulfodyne^®^-treated Calu-3 cells (Fig. 5A).

**Fig.5.**
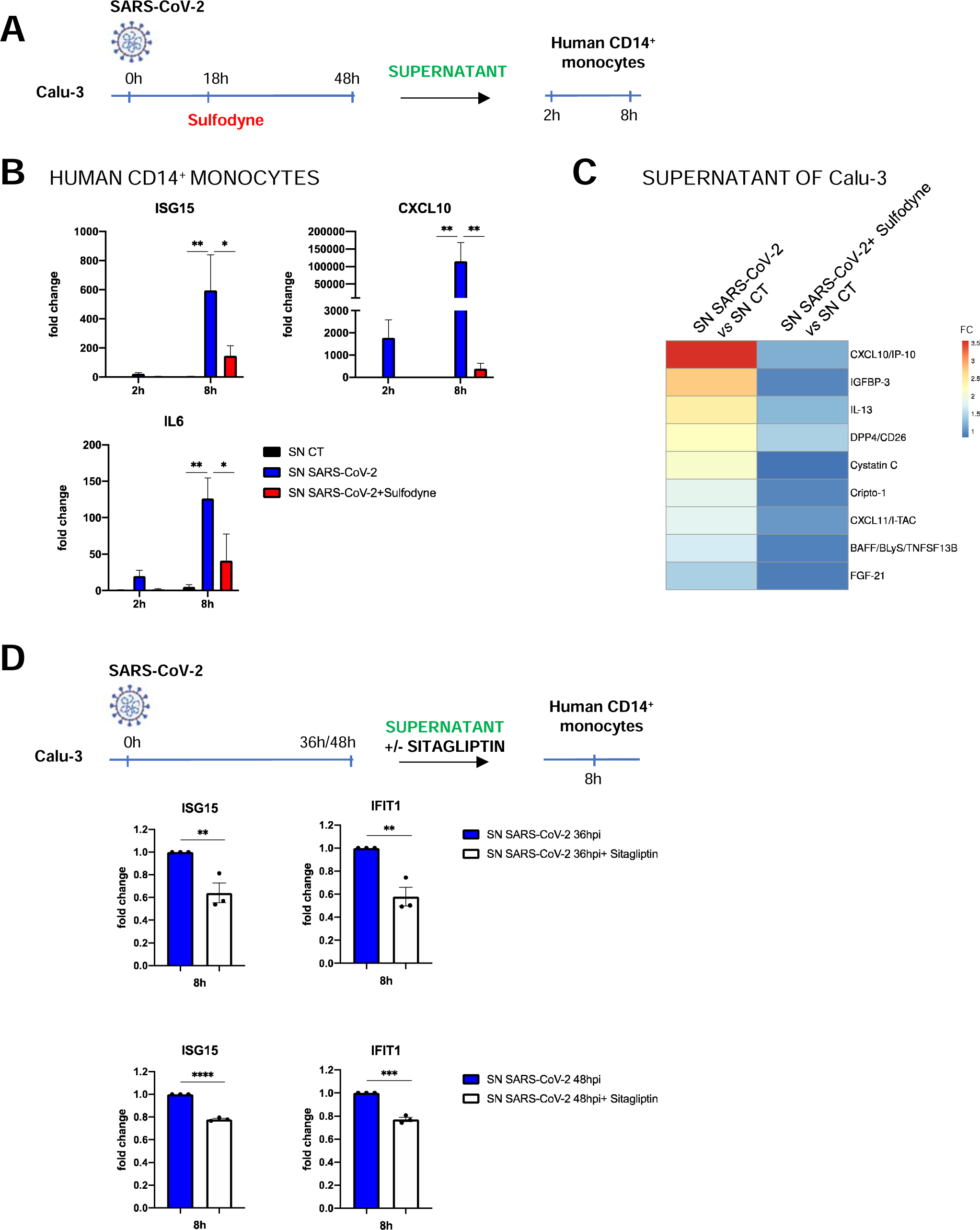
By-stander actions of Sulfodyne^®^ on human monocytes A) Schematic of experimental design. Calu-3 cells were infected with SARS-CoV-2 and treated with Sulfodyne^®^ at 18hpi. Cellular supernatants (SN) from uninfected (SN CT), SARS-CoV-2 infected (SN SARS-CoV-2) and SARS-COV-2 infected and Sulfodyne^®^ treated (SN SARS-CoV-2+Sulfodyne^®^) were collected 48hpi and used to treat human CD14^+^ monocytes. B) mRNA levels of the indicated genes were measured in human CD14^+^ monocytes 2h and 8h after exposure of indicated SN. Data are presented relative to human CD14^+^ monocytes treated with SN CT for 2h. Mean ± sem; n=3. Two-way ANOVA with Tukey’s multiple comparisons test. C) Heatmap of soluble mediators in the cellular supernatants of SARS-CoV-2 and SARS-CoV-2+Sulfodyne^®^ depicted in (A). Data are presented as fold change relative to the SN CT. D) Human CD14^+^ monocytes were exposed to SN from SARS-CoV-2 infected Calu-3 cells for 36 or 48h, in presence of Sitagliptin, a DPP4 inhibitor. mRNA levels of the indicated genes were measured 8h later and compared to mock-treated monocytes exposed to SN from infected Calu-3 cells. Mean ± sem, n=3 independent SN used on 2 different donors. Unpaired t test. * p ≤ 0.05; ** p≤ 0.01; *** p ≤ 0.001; **** p≤ 0.0001.

Exposure of human CD14^+^ monocytes to supernatant of SARS-CoV-2 infected Calu-3 cells increased mRNA levels of ISG15, CXCL10 and IL6 and this increase was significantly reduced when supernatant from Sulfodyne^®^-treated Calu-3 cells was used (Fig. 5B).

Proteome profiling of Calu-3 supernatants used to treat human monocytes showed increased protein levels of inflammatory mediators in the supernatant of infected Calu-3 cells compared to uninfected Calu-3 cells, including several known markers of severe COVID-19 such as the monocyte-recruiting chemokine CXCL10 and DPP4[26, 27], which were decreased in supernatant of Sulfodyne^®^-treated cells (Fig. 5C). DPP4 has a role in glucose metabolism[28] but increasing evidences suggest that DPP4 also acts on innate immune responses and inflammation[29]. To characterize the role of DPP4 produced by SARS-CoV-2-infected Calu-3 cells on monocytes, these were treated with Sitagliptin, a specific inhibitor of DPP4, during exposure to supernatant from SARS-CoV-2-infected Calu-3 cells (Fig. 5D, upper scheme). Sitagliptin partially reduced induction of ISG15 and IFIT1 gene expression in stimulated monocytes (Fig. 5D), suggesting a contribution of secreted DPP4 in monocyte activation and pinpointing how Sulfodyne^®^ could decrease the inflammatory response associated with SARS-CoV-2 infection.

### Sulfodyne^®^ treatment confers protection against SARS-CoV-2 infection *in vivo*

We investigated the *in vivo* effects of Sulfodyne^®^ in the Syrian golden hamsters animal model of SARS-CoV-2 infection[30]. Hamsters were treated with Sulfodyne^®^ (600mg/kg) 8 hours after intranasal inoculation of SARS-CoV-2 and twice a day for 3 days (Fig. S5A). Whereas Sulfodyne^®^ treatment did not decrease viral load in the lungs at 3dpi (Fig. S5B), it prevented loss of body weight after SARS-CoV-2 infection (Fig. S5C), suggesting a protective role of Sulfodyne^®^ against the pathological effects due to SARS-CoV-2 infection.

We then studied the effects of Sulfodyne^®^ in transgenic mice expressing the human ACE2 under the control of Keratin18 promoter (K18-hACE2) that provide a reliable mouse model for SARS-CoV-2 infection and associated severe COVID-19 disease[31]. K18-hACE2 mice were infected intranasally with Delta variant of SARS-CoV-2 and treated with Sulfodyne^®^ (600mg/kg) twice a day, starting one day before infection and for up to 14 days after infection (Fig. 6A). Sulfodyne^®^ treatment did not affect SARS-CoV-2 replication in nasal turbinates (Fig. S5D). However, Sulfodyne^®^ treatment resulted in a significant reduction in lung viral load at 3dpi, but not at 6dpi, suggesting that Sulfodyne^®^ reduced early SARS-CoV-2 replication in lung (Fig. 6B). Infection of K18-hACE2 mice showed a severe course of disease with high lethality rate at 7dpi and Sulfodyne^®^ treatment did not improve survival but could delay median survival by one day (Fig. S5E). Nevertheless, Sulfodyne^®^ treatment significantly reduced body weight loss (Fig. 6C) and decreased the early clinical symptoms (Fig. 6D, day 4 and 5). Furthermore, although in Sulfodyne^®^-treated mice the lung viral load was increased at 6dpi compared to vehicle-treated mice (Fig. 6B), no significant increase of clinical symptoms was observed at 6 and 7dpi (Fig. 6D). In line with this data, Sulfodyne^®^-treated mice displayed less severe bronchial inflammation scores at 6dpi compared to vehicle-treated animals (Fig. 6E). Altogether, these results pinpoint the beneficial role of Sulfodyne^®^ treatment *in vivo* for reducing the severity of disease associated with SARS-CoV-2 infection.

**Fig.6.**
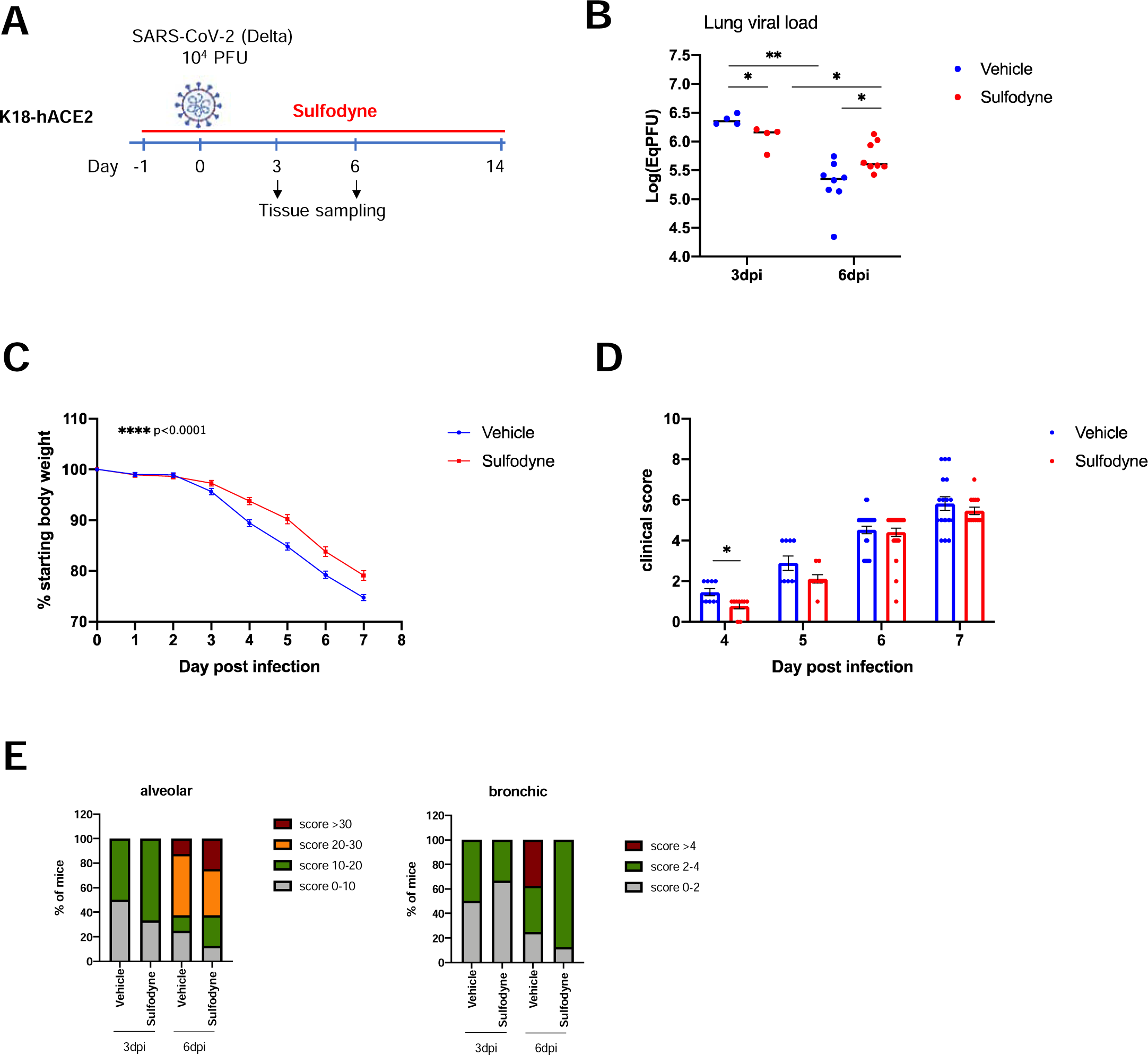
*In vivo* actions of Sulfodyne^®^ on SARS-CoV-2 symptomatology A) Schematic of experimental design. K18-hACE2 mice were intraperitoneally injected with Sulfodyne^®^ (600mg/kg) or vehicle (PBS 2%EtOH) one day prior to intranasal infection with 10^4^ PFU of SARS-CoV-2 Delta variant. Sulfodyne^®^ or vehicle were then administrated twice a day through the end of the study. B) Viral load quantification in homogenized lung tissue at 3 and 6dpi in vehicle and Sulfodyne^®^-treated mice. Individual values for each mouse and median are presented. n=4 at 3dpi; n=8 at 6dpi. Two-tailed Mann-Whitney U test. C) Body weight change during the course of infection presented as percent change compared to weight measured just before inoculation with SARS-CoV-2. Mean ± sem, n=29. Mixed-effects model comparing Sulfodyne^®^ vs vehicle treated mice. D) Clinical score was assessed for ruffled fur, hunched posture, reduced locomotion and difficult breathing in a score ranging for 0 to 2. The cumulative clinical score is indicated. Mean ± sem, n= 9 to 25. Two-tailed Mann-Whitney U test. E) Alveolar (left) and bronchic (right) histopathological score distribution in vehicle and Sulfodyne^®^-treated mice at 3 and 6dpi. n=4 (vehicle) or 3 (Sulfodyne^®^) at 3dpi; n= 8 at 6dpi.

## Discussion

In this report, we identify the natural compound Sulfodyne^®^, a stabilized form of NRF2 agonist SFN, as a potent agent against SARS-CoV-2 pathogenesis and provide evidence for multiple mechanisms that underlie the Sulfodyne^®^ beneficial effects.

As SARS-CoV-2 infection interferes with the metabolism and redox function of cellular glutathione[32], it has been suggested that the NRF2 pathway is targeted by SARS-CoV-2, leading to reduction of NRF2-mediated antioxidant responses. Indeed, in accordance with a recent study[11], we showed that NRF2 deficiency in mice infected with SARS-CoV-2 was associated with increased lung viral load and signs of exacerbated symptomatology. These results indicate a role of NRF2 in the clinical disease associated with SARS-CoV-2 infection. To characterize the role of NRF2 during SARS-CoV-2 infection, we performed transcriptomic analyses that did not evidence modulation of expression of genes linked to the NRF2 pathway in SARS-CoV-2 infected Calu-3 cells 36 hours after infection. This result extends a previous study that reported decreased NRF2 protein levels at 48 hours but not 24 hours after infection of Calu-3 cells[11] and is in accordance with the repression of the NRF2 pathway in COVID-19 patient lung biopsies[7]. Altogether, these results suggest that decreased levels of NRF2 is a late event during SARS-CoV-2 infection.

NRF2 agonists, like DMF, CDDO and SFN, possess antioxidant and cytoprotective activities, primary mediated by NRF2 activation. Their anti-inflammatory effects are not limited to the canonical activation of NRF2 but are also due to direct inhibition of nuclear factor kappa B (NF-κB), NF-κB signalling components and/or STAT3/5 activation[33]. Several reports have shown the effects of NRF2-agonists DMF and SFN during SARS-CoV-2 infection both *in vitro* and *in vivo*[7, 14]. However, the real contribution of NRF2 in the action of these agonists and the mechanisms of action of these NRF2-agonists remain elusive. Our study characterizes the signalling pathways that underlie the antiviral activity of Sulfodyne^®^ and provides essential information regarding its immunomodulatory mechanisms.

Sulfodyne^®^ treatment of epithelial cell lines elicited a more efficient inhibition of SARS-CoV-2 replication than DMF and CDDO. In these cell lines, the inhibition of SARS-CoV-2 replication by Sulfodyne^®^ occurs at post-entry stages. Although Sulfodyne^®^ treatment resulted in a strong activation of the NRF2 pathway, decreased SARS-CoV-2 replication by Sulfodyne^®^ is not dependent of NRF2 expression but associated with regulation of metabolic host pathways activated during SARS-CoV-2 infection. SARS-CoV-2 replication is structurally and functionally associated with the ER as SARS-CoV-2 ORF8 escapes degradation by host cells and induces ER stress through targeting ER chaperones and key UPR components[34]. Sulfodyne^®^ inhibited the ER stress and this inhibition may be part of the effect of Sulfodyne^®^ on SARS-CoV-2 replication. Sulfodyne^®^ can also inhibit the mTOR signalling pathway that is activated during SARS-CoV-2 infection for viral proteins translation[35]. In addition to these effects of Sulfodyne^®^ on SARS-CoV-2 replication, Sulfodyne^®^ increased expression of NRF2-target genes that activate the glutathione antioxidant system, important to restore cellular homeostatic processes during the inhibition of viral replication. Thus, our study pinpoints the beneficial role of Sulfodyne^®^ in targeting multiples host pathways essential for SARS-CoV-2 replication and in restoring the cell homeostasis.

Type I IFNs are essential in the defense against SARS-CoV-2 infection, as genetic deficiencies in IFN signalling or presence of autoantibodies neutralizing type I IFNs are strong risk factors for life-threatening COVID-19 pneumonia[36–38]. However, although rapid induction of type I IFNs limits virus propagation, late onset and continuous high levels of type I IFNs drive the immunopathology in the late phase of SARS-CoV-2 infection and are associated with poor clinical outcome[39–43]. These observations highlight the potential of immunomodulators in reducing SARS-CoV-2-driven inflammatory disease in the late stages of infection. NRF2 agonists DMF and 4-OI, by suppressing type I IFN signalling, can inhibit inflammation-associated coagulation in a model of SARS-CoV-2 infection[44]. Here, we show that Sulfodyne^®^ inhibited induction of IFNB1 and type I IFN-stimulated genes in SARS-CoV-2-infected Calu-3. Inhibition of type I IFN signalling by Sulfodyne^®^ was not restricted to epithelial cells as Sulfodyne^®^ also decreased IFN-stimulated genes, in the absence of viral replication, in both PBMC infected directly with SARS-CoV-2 and in PBMC activated with proinflammatory stimuli prior to infection with SARS-CoV-2. These results identify a role of Sulfodyne^®^ in the inhibition of type I IFN response and, together with a decreased symptomatology observed *in vivo* in both hamsters and K18-hACE2 mice, argue for a protective effect of Sulfodyne^®^ in improving the pathogenic consequences that could be associated with sustained levels of type I IFNs during SARS-CoV-2 infection. Therefore, as the IFN response is tightly regulated during SARS-CoV-2 infection, we suggest that the excessive content of interferons in patients at late stages of infection might be a biomarker for the use of Sulfodyne^®^ in clinic. Furthermore, as type I IFN production remained persistently high 8 months after SARS-CoV-2 infection in patients with long COVID[45], Sulfodyne^®^ may also be indicated in the treatment of long COVID.

Epithelial-immune crosstalk may govern many aspects of the local immune response to SARS-CoV-2 infection, including chemokines/cytokines production and inflammasome activation[24, 46], providing possible origins for the inflammatory perturbations that can occur in the lung during SARS-CoV-2 infection. SARS-CoV-2-infected epithelial cells elicited a specific proinflammatory signature in monocytes[47], which may explain the severity of COVID-19. By modelling *in vitro* the crosstalk between infected epithelial cells and monocytes, we have studied the role of Sulfodyne^®^ on the earliest events that underlie subsequent inflammatory response in bystander monocytes. We showed that SARS-CoV-2 infection in Calu-3 cells creates a pro-inflammatory microenvironment that drives innate immune responses in monocytes and that this activation could be prevented by treating epithelial cells with Sulfodyne^®^. Two known markers of severe COVID-19 were identified among the soluble mediators modulated during Sulfodyne^®^ treatment, DPP4 and CXCL10. DPP4 can act as an immunomodulator but its role during SARS-CoV-2 infection was not characterized. We showed that it might directly activate type I IFN responses in monocytes during SARS-CoV-2 infection. CXCL10 increases chemoattraction and recruitment of circulating monocytes/macrophages in tissues. Therefore, decreased levels of secreted CXCL10 found in supernatant from Sulfodyne^®^-treated Calu-3 indicated that Sulfodyne^®^ might prevent excessive monocyte infiltration. Thus, our *in vivo* and *in vitro* data show an immunomodulatory action of Sulfodyne^®^ in reducing the severity of disease associated with SARS-CoV-2 infection and are consistent with decreased activation and recruitment of myeloid cells to the lungs observed in mice treated with SFN[14].

SFN, including the encapsulated form of Sulfodyne^®^, Prostaphane[48], is orally available and well-tolerated without significant side effects and several clinical trials showed its benefits in different diseases, including lung and inflammatory diseases[33]. In rats, SFN is rapidly absorbed, reaches a maximum plasma concentration 4 hours after absorption and has a half-life around 2 hours[49]. In human, the bioavailability of SFN is variable and depends on concentration and type of formulations administered[50]. The discrepancy between the *in vitro* and *in vivo* effects of Sulfodyne^®^ on SARS-CoV-2 replication might be related to its bioavailability. Thus, further studies in human to characterize Sulfodyne^®^ pharmacokinetic properties are required before any use of Sulfodyne^®^ to treat the pathogenic consequences of SARS-CoV-2 infection.

In conclusion, by acting directly on immune cells and by decreasing the innate immune response and the secreted signals produced by SARS-CoV-2-infected epithelial cells and the subsequent recruitment and activation of monocytes, Sulfodyne^®^ contributes to reduce pathological effects associated with sustained activation of immune cells during SARS-CoV-2 infection in lungs. Considering its beneficial role in conferring protection during SARS-CoV-2 infection at multiples levels, Sulfodyne^®^ appears as a promising therapeutic agent of COVID-19 symptomatology.

## Materials and Methods

### Cell culture, treatments and virus

Human airway epithelial cells Calu-3 (ATCC) were cultured in EMEM supplemented with 15% FCS. Human colorectal adenocarcinoma cells Caco-2 (ATCC) were cultured in DMEM supplemented with 10% FCS and 1% not essential amino-acids.

High-purity Sulfodyne^®^ (https://ingoodbyolga.com/en/ingredient/sulfodyne/), an health ingredient of broccoli seeds extract titrated in 5% of natural, active and stablized SFN, was provided by Ingood by Olga company, dissolved in PBS and used, unless otherwise indicated, at 50ug/ml, a concentration equivalent to 14uM of SFN. DMF (Sigma-Aldrich) was used at 150μM, CDDO-Imidazolide (Sigma-Aldrich) at 50nM, Rapamycin (Sigma-Aldrich) at 1μM, Sitagliptin (Selleckchem) at 100μM, LPS (Sigma-Aldrich) at 100 ng/ml. MTT cytotoxicity assay (CliniSciences) was performed according to the manufacturer’s instructions.

SARS-CoV-2 Wuhan, Beta (B.1.351 strain) and Delta (B.1.617.2 strain) strains were provided by the National Reference Center for Respiratory Viruses (Institut Pasteur, Paris, France). All procedures involving infectious SARS-CoV-2 were performed in biosafety level 3 (BSL-3) facilities at IDMIT (CEA, Fontenay-aux-Roses, France).

### RNA extraction and RT-qPCR

Total cellular RNA was extracted with the nucleospin RNA plus XS (Macherey-Nagel). Viral RNA in supernatant cell culture was isolated with the Nucleospin Dx virus (Macherey-Nagel). Reverse transcription was performed with random primers and Superscript IV (Life Technologies) and quantitative PCR with the Power SYBR green PCR master mix (Applied Biosystems) for host transcripts or TaqMan™ Fast Advanced Master Mix (Applied Biosystems) for viral transcripts. The relative quantification of mRNA levels was calculated using the threshold cycle (2−ΔΔCT) method with ACTB RNA as an internal control.

The following primer pairs were used (forward and reverse):

HMOX1: CCAGGCAGAGAATGCTGAGTTC and AAGACTGGGCTCTCCTTGTTGC

TXNRD1: GTTACTTGGGCATCCCTGGTGA and CGCACTCCAAAGCGACATAGGA

HSPA5: CTGTCCAGGCTGGTGTGCTCT and CTTGGTAGGCACCACTGTGTTC

DDIT3: GGTATGAGGACCTGCAAGAGGT and CTTGTGACCTCTGCTGGTTCTG

IFNB1: TCATGAGTTTTCCCCTGGTG and GTTGAGAACCTCCTGGCTAATG

IFIT1: GCCTTGCTGAAGTGTGGAGGAA and ATCCAGGCGATAGGCAGAGATC

ISG15: CTCTGAGCATCCTGGTGAGGAA and AAGGTCAGCCAGAACAGGTCGT

CXCL10: CGCTGTACCTGCATCAGCATTAG and CTGGATTCAGACATCTCTTCTCACC

IL6: TCCAGAACAGATTTGAGAGTAGTG and GCATTTGTGGTTGGGTCAGG

ACTB: CACCATTGGCAATGAGCGGTTC and AGGTCTTTGCGGATGTCCACGT

IP4 (SARS-CoV-2 genomic): GGTAACTGGTATGATTTCG and CTGGTCAAGGTTAATATAGG, (probe) TCATACAAACCACGCCAGG;

E gene/Leader (SARS-CoV-2 subgenomic): CGATCTCTTGTAGATCTGTTCTC and ATATTGCAGCAGTACGCACACA, (probe) ACACTAGCCATCCTTACTGCGCTTCG.

### RNA-seq

Calu-3 cells were infected with SARS-CoV-2 and treated or not with Sulfodyne^®^ at 12hpi. Cells were harvested at 0hpi (mock), 12hpi, 18hpi and 36hpi for RNA-seq. Four independent biological replicates were performed per experimental condition. Library construction, sequencing on Oxford Nanopore Technologies sequencer and bioinformatics analysis were performed by Life&Soft company. A total of 100 ng of RNA per sample was retrotranscribed by a strand-switching technique using Maxima H Minus Reverse Transcriptase (ThermoFisher) to synthesize a doublestranded cDNA. PCR, barcode and adapter attachment were performed according to PCR-cDNA Barcoding kit (Oxford Nanopore Technologies). Samples were quantified using QuBit dsDNA HS kit (ThermoFisher) before loading on R9.4.1 Flow cells using the GridION instrument (Minknow version: 21.11.6). Sequence reads were converted into FASTQ files. Reads under 300 bp or with a quality score under 9 were discarded. The remaining reads were mapped on the human GRCh38.p13 and SARS-CoV-2 ASM985889v3 transcriptome of reference using minimap2 version 2.17[51]. To quantify transcripts, the resulting alignments were given to Salmon version 1.6.0[52].Differentially expressed genes (DEG) were determined using DESeq2. Enrichment analysis was performed using Metascape or GSEA.

### Protein extraction and capillary-based immunoblotting (WES Simple western)

Total protein extraction was performed with RIPA buffer (20mM HEPES pH 7.6, 150mM NaCl, 1% NP4, 0.25% sodium deoxycholate, 10% glycerol). For nuclear extracts, cells were resuspended in hypotonic buffer (10mM HEPES pH 7.6, 1.5mM MgCl_2_, 10mM KCl and protease inhibitor cocktail) and nuclei were extracted in 20mM HEPES pH 7.6, 20% glycerol, 420 mM NaCl, 1.5 mM MgCl_2_, 0.2 mM EDTA, 0.5mM DTT and protease inhibitor cocktail.

Proteins were run on the WES Simple western system using a 12-230 kDa separation module (Bio-Techne).

### Antibodies

Antibodies used for capillary-based immunoblotting are: Phospho-p70 S6 Kinase (Thr389; Cell Signaling 9205S), p70 S6 Kinase (Cell Signaling 2708S), Phospho-S6 Ribosomal Protein (Ser235/236; Cell Signaling 4858S), NRF2 (Cell Signaling 12721), GAPDH (Cell Signaling 2118), HDAC1 (Abcam ab7028). ACE2 antibody (R&D AF933) was used for Flow Cytometry. SARS-CoV-2 Spike antibody (Active Motif, clone AM001414) was used for neutralization at 10nM.

### ShRNA

For NRF2 knock-down in Calu-3 and Caco2 cells, the sequence of the shRNA targeting 3UTR of human NRF2 gene was obtained from Sigma Aldrich (TRCN0000007555; target sequence: GCTCCTACTGTGATGTGAAAT) and cloned in the pTRIP-MND-GFP-H1 lentiviral vector. After lentiviral production and cells transduction, GFP-positive cells were isolated by fluorescence-activated cell sorting and used in functional assays.

### Cytokine measurement

Soluble mediators were detected in Calu-3 culture supernatant by the Proteome Profiler Human XL Cytokine Array Kit (R&D Biosystems) according to the manufacturer’s instructions.

### Primary cells

Peripheral blood was obtained from healthy donors in accordance to the ethical guidelines at CEA (Fontenay-aux-Roses, France) and Peripheral Blood Mononuclear Cells (PBMC) were isolated by Ficoll (Eurobio Scientific) gradient centrifugation. Monocytes were isolated from PBMC using CD14 microbeads (Miltenyi Biotec) according to the manufacturer’s instructions. PBMC and CD14+ monocytes were cultured in RPMI supplemented with 10% FCS.

### Hamsters and *in vivo* infection

Animal housing and experimental procedures were conducted by Oncodesign, according to the French and European Regulations and the Institutional Animal Care and Use Committee of CEA approved by French authorities (CETEA DSV-n°44). The animal BSL3 facility is authorized by the French authorities (Agreement N° D92-032-02).

6-8 weeks old female Syrian golden hamsters were infected with 10^4^ pfu TCID_50_ of SARS-CoV-2 (Slovakia/SK-BMC5/2020, GISAID EPI_ISL_417879) by intranasal route under a total volume of 70 µL (35 µL per nostril) on Isoflurane-anesthetized animals. Sulfodyne^®^ (600mg/kg) was administered orally at 8hpi and then twice daily with an 8h interval between each delivery on one given day. Body weight was monitored daily. Animals were euthanized at 3dpi and the right lung lobe collected for RNA extraction (Macherey Nagel NucleoSpin) and viral load quantification by qRT-PCR using the IP4 set of primers and probe.

### Mice and *in vivo* infection

B6.129P2-*Nfe2l2^tm1Mym^* (*Nrf2* KO in the text) and C57BL/6J mice were bred at the TAAM (https://www.taam.cnrs.fr/) under SPF conditions. B6.Cg-Tg(K18-ACE2)2Prlmn/J mice (stock #034860, K18-hACE2 in the text) were imported from The Jackson Laboratory (Bar Harbor, ME, USA) and bred at the Institut Pasteur under strict SPF conditions. Infection studies were performed in animal BSL-3 facility at the Institut Pasteur. All animal work was approved by the Institut Pasteur Ethics Committee (project dap 200008) and authorized by the French Ministry of Research under project 24613 in compliance with the European and French regulations.

Infection experiments were performed on male and female mice, aged 7-12 weeks (K18-hACE2) or 7-20 weeks (*Nrf2* KO). Groups were composed to be sex- and age-matched. Anesthetized (ketamine/xylazine) mice were inoculated intranasally with 6 x10^4^ plaque-forming units (PFU) of SARS-CoV-2 Beta variant (B.1.351 strain, GISAID ID: EPI_ISL_964916) in a 24µl volume (*Nrf2* KO) or 10^4^ PFU of SARS-CoV-2 Delta variant (B.1.617.2 strain, GISAID ID: EPI_ISL_3030060) in a 40µl volume (K18-hACE2). For K18-hACE2 mice experiments, Sulfodyne^®^ (600mg/kg) was administered by intraperitoneal injection two times per day, starting one day before infection and for up to 14 days after infection. One group of mice was followed for 14 days post-infection (dpi), for clinical signs and body weight. Each of four criteria (ruffled fur, hunched posture, reduced locomotion and difficult breathing) received a 0-2 score and were added into a global score. Mice were euthanized when reaching 25% body weight loss or a global score of 8. Other groups of mice were euthanized at specific dpi for tissue collection.

### Viral load

Groups of mice were euthanized at 3 or 6 dpi for measurement of viral load and viral titer in lung homogenates and nasal turbinates homogenates. Right lung lobe and nasal turbinates were dissected and frozen at −80°C. Samples were homogenized in 400 μl of cold PBS using lysing matrix M (MP Biomedical) and a MP Biomedical FastPrep 24 Tissue Homogenizer. Viral RNA was extracted in PBS using an extraction robot IDEAL-32 (IDsolutions) and the NucleoMag Pathogen extraction kit (Macherey Nagel). Viral RNA quantification was performed by RT-qPCR using the IP4 set of primers and probe and the Luna Universal Probe One-Step RT-qPCR Kit (NEB). Serial dilutions of a titrated viral stock were analyzed simultaneously to express viral loads as eqPFU per gram of tissue.

### Histopathological analysis

The left lung lobe was fixed in 10% phosphate buffered formalin for 7 days. Histological analysis was performed on paraffin-embedded sections and stained with hematoxylin-eosin. The alveolar scoring system considered alveolar thickness, interstitial inflammation, alveolar inflammation and alveolar inflammation at the surface. The bronchi scoring system considered inflammation and activation.

### Statistical analysis

Data were analyzed using GraphPad Prism 9. Specific statistical tests are detailed in the figure legends. Statistical significance: * p ≤ 0.05; ** p≤ 0.01; *** p ≤ 0.001; **** p≤ 0.0001.

### Data availability

The RNAseq data are available in the GEO database under accession code GSE243268.

## Acknowledgments

We are grateful for the excellent contributions of the staff at the IDMIT center. We thank Sylvie van Der Werf at the National Reference Centre for Respiratory Viruses (Institut Pasteur) for providing the SARS-CoV-2 virus; Coralie Perrigault, Nicolas Plu and Théodoros Efstathiou at the Olga company for providing Sulfodyne^®^; Eric Ginoux, Léo d’Agata, Ségolène Diry, Natalia Nuñez and Cassandra Gaspar at the Life&Soft company for excellent assistance with RNAseq experiments; Nathalie Gault, Vilma Barroca, Cécile Ducrot and Claire Torres for helpful discussions and technical support; Didier Busso and Guillame Piton at the CIGEX Platform (iRCM/CEA) for lentiviral productions and the staff of Oncodesign for hamster experiments.

## Author contributions

PHR, VP and FF conceived and designed the study, analyzed data, supervised the project and wrote the manuscript; LC, XM, VP and FF conducted experiments and data analyses; SM and QP performed histological analysis; SGM, AB and JBS assisted with preliminary experiments; NDB, XM and RLG provided resources, scientific expertise and intellectual support. The authors read and approved the final manuscript.

## Supporting Figure Legends

Fig.S1

A) Schematic of experimental design. WT and *Nrf2* KO mice were intranasally infected with SARS-CoV-2 Beta variant (6×10^4^ PFU).

B) Viral RNA quantification in homogenized lung tissue at 6dpi in WT and *Nrf2* KO mice. Individual values for each mouse and median are presented, n=13. Two-tailed Mann-Whitney U test.

C) Percent starting body weight. Mean ± sem, n=17 to 30. Mixed-effects model test comparing *Nrf2* KO vs WT.

D) Clinical score was assessed for ruffled fur, hunched posture, reduced locomotion and difficult breathing in a score ranging for 0 to 2. The cumulative clinical score is indicated. WT mice did not exhibit any clinical signs. Mean ± sem, n=13.

E) Representative haematoxylin-eosin staining (left) and histopathological score distribution (right) of the lungs of WT and *Nrf2* KO mice at 6dpi. n=12 (WT) or 13 (Nrf2 KO).

Fig. S2

A) Representative WES Simple assay for NRF2, HDAC1 and GAPDH in nuclear and cytoplasmic extracts from Calu-3 cells treated with Sulfodyne^®^ (50ug/ml) for 6h.

B) Cell viability of Calu-3 cells treated with Sulfodyne^®^ (50ug/ml) for 6 and 30h, measured by MTT assay. Data are presented as percent of untreated cells. Mean ± sem, n=3.

C) NRF2-target gene *HMOX1* mRNA levels in Calu-3 infected and treated with Sulfodyne^®^ at indicated times before or after infection. Dare are shown as fold change over untreated (UT) cells at 36hpi. Mean ± sem, n=3.

Fig. S3

A) Validation of anti-Spike neutralizing antibody (Spike Ab). SARS-CoV-2 was pre-incubated in presence or not of the Spike Ab for 1hour at 37°C before infection of Calu-3 cells. Genomic and sub-genomic viral RNA were measured 36hpi. Mean ± sem, n=2.

Fig. S4

A) Heatmap of normalized expression levels of DEG (up and down-regulated) in SARS-CoV-2 Calu-3 at 36hpi vs mock.

B) Metascape pathway analysis comparing DEG during infection (COV2_36hpi vs mock) and DEG by Sulfodyne^®^ treatment (COV2_Sulfodyne^®^_36hpi vs COV2_36hpi). Circle size indicates statistically significance of the enrichment.

C-D) Representative WES Simple assay for NRF2 and HDAC1 in nuclear extract from Calu-3 (C) or Caco2 (D) transduced with shRNA targeting NRF2 (shNRF2) or a non-targeting control (shCT) and treated with Sulfodyne^®^ for 6h.

E) Viral genomic and NRF2-target gene *TXNRD1* mRNA levels in shCT and shNRF2 Caco2 infected with SARS-CoV-2 for 48hpi. Cells were treated with Sulfodyne^®^ at 18hpi. Data are presented as fold change over ShCT UT cells at 48hpi for genomic RNA and as fold change over uninfected cells (0hpi) for *TXNRD1.* Mean ± sem, n=3. Two-way ANOVA with Sidak’s multiple comparisons test vs shCT.

F) WES Simple assay for mTOR substrate proteins and GAPDH in Caco2 cells infected for 48h and treated with Sulfodyne^®^ or Rapamycin at 18hpi.

Fig. S5

A) Schematic of experimental design. Syrian Golden hamsters (females, 6-8 weeks old) were intranasally infected with SARS-CoV-2 (10^4^ PFU) 8h before oral administration of Sulfodyne^®^ (600mg/kg) or vehicle.

B) Lung viral load at 3dpi in vehicle and Sulfodyne^®^-treated hamsters. Mean ± sem, n=6. Two-tailed Mann-Whitney U test.

C) Percent of starting body weight in vehicle and Sulfodyne^®^-treated hamsters. Mean ± sem, n=6. Two-way ANOVA with Sidak’s multiple comparisons test comparing Sulfodyne^®^ vs vehicle.

D) Viral RNA quantification in nasal turbinates at 3 and 6dpi in vehicle and Sulfodyne^®^-treated K18-hACE2 mice. Individual values for each mouse and median are presented, n=4 at 3dpi and n=8 at 6dpi. Two-tailed Mann-Whitney U test.

E) Kaplan–Meier plot of survival of vehicle and Sulfodyne^®^-treated K18-hACE2 mice. n=17. ns: not significant,* p ≤ 0.05; ** p≤ 0.01

## References

1. Huang C, Wang Y, Li X, Ren L, Zhao J, Hu Y, et al. Clinical features of patients infected with 2019 novel coronavirus in Wuhan, China. Lancet. 2020;395: 497–506. doi:10.1016/S0140-6736(20)30183-5

2. Toussi SS, Hammond JL, Gerstenberger BS, Anderson AS. Therapeutics for COVID-19. Nat Microbiol. 2023;8: 771–786. doi:10.1038/s41564-023-01356-4

3. Tonelli C, Chio IIC, Tuveson DA. Transcriptional Regulation by Nrf2. Antioxid Redox Signal. 2018;29: 1727–1745. doi:10.1089/ars.2017.7342

4. Yamamoto M, Kensler TW, Motohashi H. The KEAP1-NRF2 System: a Thiol-Based Sensor-Effector Apparatus for Maintaining Redox Homeostasis. Physiol Rev. 2018;98: 1169– 1203. doi:10.1152/physrev.00023.2017

5. Kobayashi EH, Suzuki T, Funayama R, Nagashima T, Hayashi M, Sekine H, et al. Nrf2 suppresses macrophage inflammatory response by blocking proinflammatory cytokine transcription. Nat Commun. 2016;7: 11624. doi:10.1038/ncomms11624

6. Ramezani A, Nahad MP, Faghihloo E. The role of Nrf2 transcription factor in viral infection. J Cell Biochem. 2018;119: 6366–6382. doi:10.1002/jcb.26897

7. Olagnier D, Farahani E, Thyrsted J, Blay-Cadanet J, Herengt A, Idorn M, et al. SARS-CoV2-mediated suppression of NRF2-signaling reveals potent antiviral and anti-inflammatory activity of 4-octyl-itaconate and dimethyl fumarate. Nat Commun. 2020;11: 4938. doi:10.1038/s41467-020-18764-3

8. Gümüş H, Erat T, Öztürk İ, Demir A, Koyuncu I. Oxidative stress and decreased Nrf2 level in pediatric patients with COVID-19. Journal of Medical Virology. 2022;94: 2259–2264. doi:10.1002/jmv.27640

9. Zhang S, Wang J, Wang L, Aliyari S, Cheng G. SARS-CoV-2 virus NSP14 Impairs NRF2/HMOX1 activation by targeting Sirtuin 1. Cell Mol Immunol. 2022;19: 872–882. doi:10.1038/s41423-022-00887-w

10. Liu L, Du J, Yang S, Zheng B, Shen J, Huang J, et al. SARS-CoV-2 ORF3a sensitizes cells to ferroptosis via Keap1-NRF2 axis. Redox Biol. 2023;63: 102752. doi:10.1016/j.redox.2023.102752

11. Qu Y, Haas de Mello A, Morris DR, Jones-Hall YL, Ivanciuc T, Sattler RA, et al. SARS-CoV-2 Inhibits NRF2-Mediated Antioxidant Responses in Airway Epithelial Cells and in the Lung of a Murine Model of Infection. Microbiol Spectr. 2023;11: e0037823. doi:10.1128/spectrum.00378-23

12. McCord JM, Hybertson BM, Cota-Gomez A, Geraci KP, Gao B. Nrf2 Activator PB125^®^ as a Potential Therapeutic Agent against COVID-19. Antioxidants (Basel). 2020;9: 518. doi:10.3390/antiox9060518

13. Rothan HA, Zhong Y, Sanborn MA, Teoh TC, Ruan J, Yusof R, et al. Small molecule grp94 inhibitors block dengue and Zika virus replication. Antiviral Res. 2019;171: 104590. doi:10.1016/j.antiviral.2019.104590

14. Ordonez AA, Bullen CK, Villabona-Rueda AF, Thompson EA, Turner ML, Merino VF, et al. Sulforaphane exhibits antiviral activity against pandemic SARS-CoV-2 and seasonal HCoV-OC43 coronaviruses in vitro and in mice. Commun Biol. 2022;5: 242. doi:10.1038/s42003-022-03189-z

15. Chen Z, Du R, Cooper L, Achi JG, Dong M, Ran Y, et al. Sulforaphane is a reversible covalent inhibitor of 3-chymotrypsin-like protease of SARS-CoV-2. J Med Virol. 2023;95: e28609. doi:10.1002/jmv.28609

16. Gasparello J, D’Aversa E, Papi C, Gambari L, Grigolo B, Borgatti M, et al. Sulforaphane inhibits the expression of interleukin-6 and interleukin-8 induced in bronchial epithelial IB3-1 cells by exposure to the SARS-CoV-2 Spike protein. Phytomedicine. 2021;87: 153583. doi:10.1016/j.phymed.2021.153583

17. Kiser C, Gonul CP, Olcum M, Genc S. Inhibitory effects of sulforaphane on NLRP3 inflammasome activation. Mol Immunol. 2021;140: 175–185. doi:10.1016/j.molimm.2021.10.014

18. Fahey JW, Wade KL, Wehage SL, Holtzclaw WD, Liu H, Talalay P, et al. Stabilized sulforaphane for clinical use: Phytochemical delivery efficiency. Mol Nutr Food Res. 2017;61. doi:10.1002/mnfr.201600766

19. Zhao S, Ghosh A, Lo C-S, Chenier I, Scholey JW, Filep JG, et al. Nrf2 Deficiency Upregulates Intrarenal Angiotensin-Converting Enzyme-2 and Angiotensin 1-7 Receptor Expression and Attenuates Hypertension and Nephropathy in Diabetic Mice. Endocrinology. 2018;159: 836–852. doi:10.1210/en.2017-00752

20. Wang Y, Ma J, Jiang Y. Transcription factor Nrf2 as a potential therapeutic target for COVID-19. Cell Stress Chaperones. 2023;28: 11–20. doi:10.1007/s12192-022-01296-8

21. Appelberg S, Gupta S, Svensson Akusjärvi S, Ambikan AT, Mikaeloff F, Saccon E, et al. Dysregulation in Akt/mTOR/HIF-1 signaling identified by proteo-transcriptomics of SARS-CoV-2 infected cells. Emerg Microbes Infect. 2020;9: 1748–1760. doi:10.1080/22221751.2020.1799723

22. Lei X, Dong X, Ma R, Wang W, Xiao X, Tian Z, et al. Activation and evasion of type I interferon responses by SARS-CoV-2. Nat Commun. 2020;11: 3810. doi:10.1038/s41467-020-17665-9

23. Li M, Ferretti M, Ying B, Descamps H, Lee E, Dittmar M, et al. Pharmacological activation of STING blocks SARS-CoV-2 infection. Sci Immunol. 2021;6: eabi9007. doi:10.1126/sciimmunol.abi9007

24. Thorne LG, Reuschl A-K, Zuliani-Alvarez L, Whelan MVX, Turner J, Noursadeghi M, et al. SARS-CoV-2 sensing by RIG-I and MDA5 links epithelial infection to macrophage inflammation. EMBO J. 2021;40: e107826. doi:10.15252/embj.2021107826

25. Kazmierski J, Friedmann K, Postmus D, Emanuel J, Fischer C, Jansen J, et al. Nonproductive exposure of PBMCs to SARS-CoV-2 induces cell-intrinsic innate immune responses. Mol Syst Biol. 2022;18: e10961. doi:10.15252/msb.202210961

26. Gudowska-Sawczuk M, Mroczko B. What Is Currently Known about the Role of CXCL10 in SARS-CoV-2 Infection? Int J Mol Sci. 2022;23: 3673. doi:10.3390/ijms23073673

27. Sebastián-Martín A, Sánchez BG, Mora-Rodríguez JM, Bort A, Díaz-Laviada I. Role of Dipeptidyl Peptidase-4 (DPP4) on COVID-19 Physiopathology. Biomedicines. 2022;10: 2026. doi:10.3390/biomedicines10082026

28. Mulvihill EE, Drucker DJ. Pharmacology, physiology, and mechanisms of action of dipeptidyl peptidase-4 inhibitors. Endocr Rev. 2014;35: 992–1019. doi:10.1210/er.2014-1035

29. Yazbeck R, Jaenisch SE, Abbott CA. Dipeptidyl peptidase 4 inhibitors: Applications in innate immunity? Biochem Pharmacol. 2021;188: 114517. doi:10.1016/j.bcp.2021.114517

30. Imai M, Iwatsuki-Horimoto K, Hatta M, Loeber S, Halfmann PJ, Nakajima N, et al. Syrian hamsters as a small animal model for SARS-CoV-2 infection and countermeasure development. Proc Natl Acad Sci U S A. 2020;117: 16587–16595. doi:10.1073/pnas.2009799117

31. Golden JW, Cline CR, Zeng X, Garrison AR, Carey BD, Mucker EM, et al. Human angiotensin-converting enzyme 2 transgenic mice infected with SARS-CoV-2 develop severe and fatal respiratory disease. JCI Insight. 2020;5: e142032, 142032. doi:10.1172/jci.insight.142032

32. Bartolini D, Stabile AM, Bastianelli S, Giustarini D, Pierucci S, Busti C, et al. SARS-CoV2 infection impairs the metabolism and redox function of cellular glutathione. Redox Biol. 2021;45: 102041. doi:10.1016/j.redox.2021.102041

33. Cuadrado A, Pajares M, Benito C, Jiménez-Villegas J, Escoll M, Fernández-Ginés R, et al. Can Activation of NRF2 Be a Strategy against COVID-19? Trends Pharmacol Sci. 2020;41: 598–610. doi:10.1016/j.tips.2020.07.003

34. Liu P, Wang X, Sun Y, Zhao H, Cheng F, Wang J, et al. SARS-CoV-2 ORF8 reshapes the ER through forming mixed disulfides with ER oxidoreductases. Redox Biol. 2022;54: 102388. doi:10.1016/j.redox.2022.102388

35. Mullen PJ, Garcia G, Purkayastha A, Matulionis N, Schmid EW, Momcilovic M, et al. SARS-CoV-2 infection rewires host cell metabolism and is potentially susceptible to mTORC1 inhibition. Nat Commun. 2021;12: 1876. doi:10.1038/s41467-021-22166-4

36. Bastard P, Rosen LB, Zhang Q, Michailidis E, Hoffmann H-H, Zhang Y, et al. Autoantibodies against type I IFNs in patients with life-threatening COVID-19. Science. 2020;370: eabd4585. doi:10.1126/science.abd4585

37. Zhang Q, Bastard P, Liu Z, Le Pen J, Moncada-Velez M, Chen J, et al. Inborn errors of type I IFN immunity in patients with life-threatening COVID-19. Science. 2020;370: eabd4570. doi:10.1126/science.abd4570

38. Bastard P, Gervais A, Le Voyer T, Rosain J, Philippot Q, Manry J, et al. Autoantibodies neutralizing type I IFNs are present in ∼4% of uninfected individuals over 70 years old and account for ∼20% of COVID-19 deaths. Sci Immunol. 2021;6: eabl4340. doi:10.1126/sciimmunol.abl4340

39. Park A, Iwasaki A. Type I and Type III Interferons - Induction, Signaling, Evasion, and Application to Combat COVID-19. Cell Host Microbe. 2020;27: 870–878. doi:10.1016/j.chom.2020.05.008

40. Lee JS, Shin E-C. The type I interferon response in COVID-19: implications for treatment. Nat Rev Immunol. 2020;20: 585–586. doi:10.1038/s41577-020-00429-3

41. Zhou Z, Ren L, Zhang L, Zhong J, Xiao Y, Jia Z, et al. Heightened Innate Immune Responses in the Respiratory Tract of COVID-19 Patients. Cell Host Microbe. 2020;27: 883–890.e2. doi:10.1016/j.chom.2020.04.017

42. Lee JS, Park S, Jeong HW, Ahn JY, Choi SJ, Lee H, et al. Immunophenotyping of COVID-19 and influenza highlights the role of type I interferons in development of severe COVID-19. Sci Immunol. 2020;5: eabd1554. doi:10.1126/sciimmunol.abd1554

43. Galani I-E, Rovina N, Lampropoulou V, Triantafyllia V, Manioudaki M, Pavlos E, et al. Untuned antiviral immunity in COVID-19 revealed by temporal type I/III interferon patterns and flu comparison. Nat Immunol. 2021;22: 32–40. doi:10.1038/s41590-020-00840-x

44. Ryan TAJ, Hooftman A, Rehill AM, Johansen MD, Brien ECO, Toller-Kawahisa JE, et al. Dimethyl fumarate and 4-octyl itaconate are anticoagulants that suppress Tissue Factor in macrophages via inhibition of Type I Interferon. Nat Commun. 2023;14: 3513. doi:10.1038/s41467-023-39174-1

45. Phetsouphanh C, Darley DR, Wilson DB, Howe A, Munier CML, Patel SK, et al. Immunological dysfunction persists for 8 months following initial mild-to-moderate SARS-CoV-2 infection. Nat Immunol. 2022;23: 210–216. doi:10.1038/s41590-021-01113-x

46. Barnett KC, Xie Y, Asakura T, Song D, Liang K, Taft-Benz SA, et al. An epithelial-immune circuit amplifies inflammasome and IL-6 responses to SARS-CoV-2. Cell Host Microbe. 2023;31: 243–259.e6. doi:10.1016/j.chom.2022.12.005

47. Leon J, Michelson DA, Olejnik J, Chowdhary K, Oh HS, Hume AJ, et al. A virus-specific monocyte inflammatory phenotype is induced by SARS-CoV-2 at the immune-epithelial interface. Proc Natl Acad Sci U S A. 2022;119: e2116853118. doi:10.1073/pnas.2116853118

48. Cipolla BG, Mandron E, Lefort JM, Coadou Y, Della Negra E, Corbel L, et al. Effect of Sulforaphane in Men with Biochemical Recurrence after Radical Prostatectomy. Cancer Prev Res (Phila). 2015;8: 712–719. doi:10.1158/1940-6207.CAPR-14-0459

49. Hu R, Hebbar V, Kim B-R, Chen C, Winnik B, Buckley B, et al. In vivo pharmacokinetics and regulation of gene expression profiles by isothiocyanate sulforaphane in the rat. J Pharmacol Exp Ther. 2004;310: 263–271. doi:10.1124/jpet.103.064261

50. Yagishita Y, Fahey JW, Dinkova-Kostova AT, Kensler TW. Broccoli or Sulforaphane: Is It the Source or Dose That Matters? Molecules. 2019;24: 3593. doi:10.3390/molecules24193593

51. Li H. Minimap2: pairwise alignment for nucleotide sequences. Bioinformatics. 2018;34: 3094–3100. doi:10.1093/bioinformatics/bty191

52. Patro R, Duggal G, Love MI, Irizarry RA, Kingsford C. Salmon provides fast and bias-aware quantification of transcript expression. Nat Methods. 2017;14: 417–419. doi:10.1038/nmeth.4197

