## Supplementary figures and images for "Multi-target mode of action of Sulfodyne^®^, a stabilized Sulforaphane, against pathogenic effects of SARS-CoV-2 infection"

### Supporting Figures

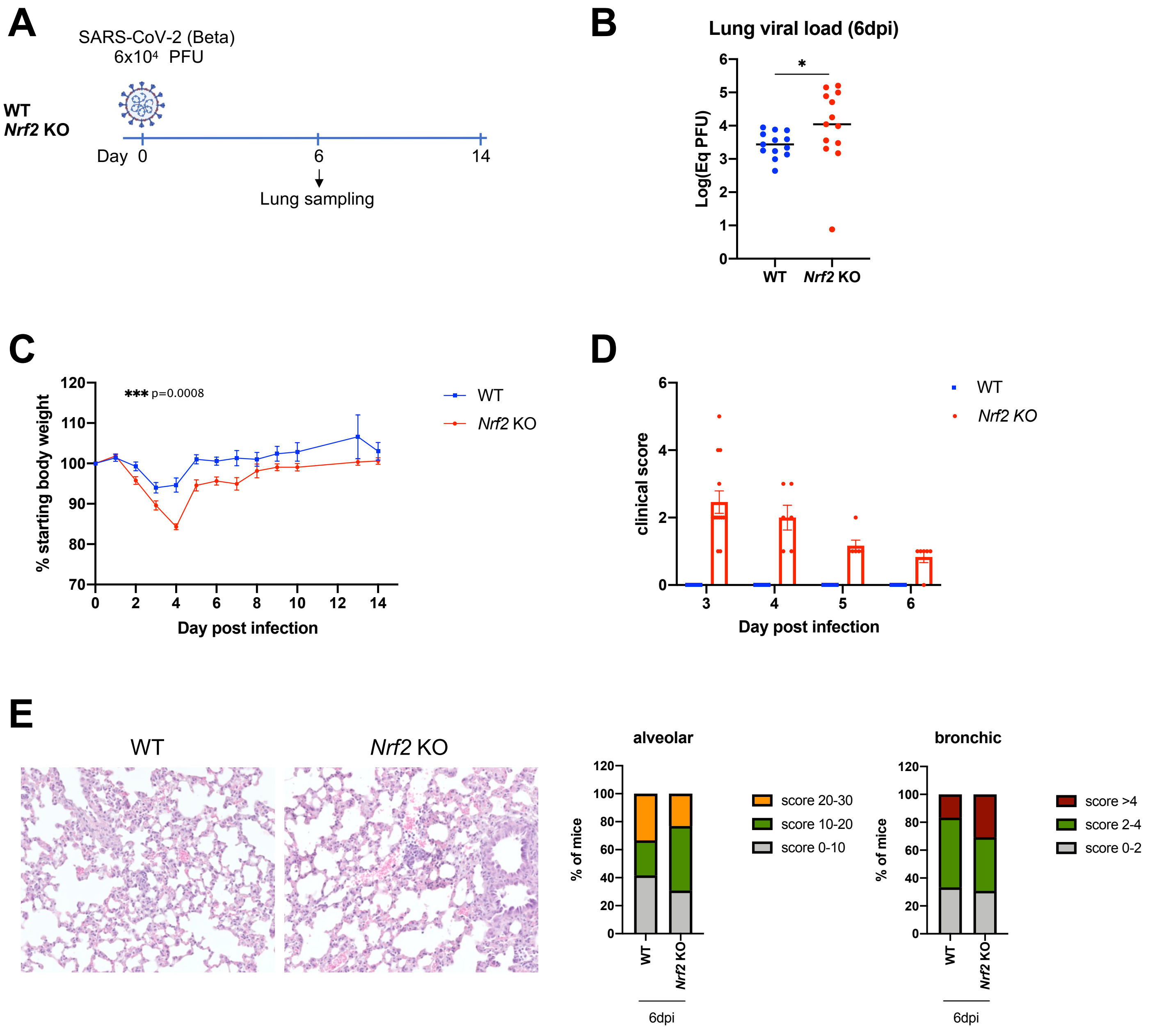

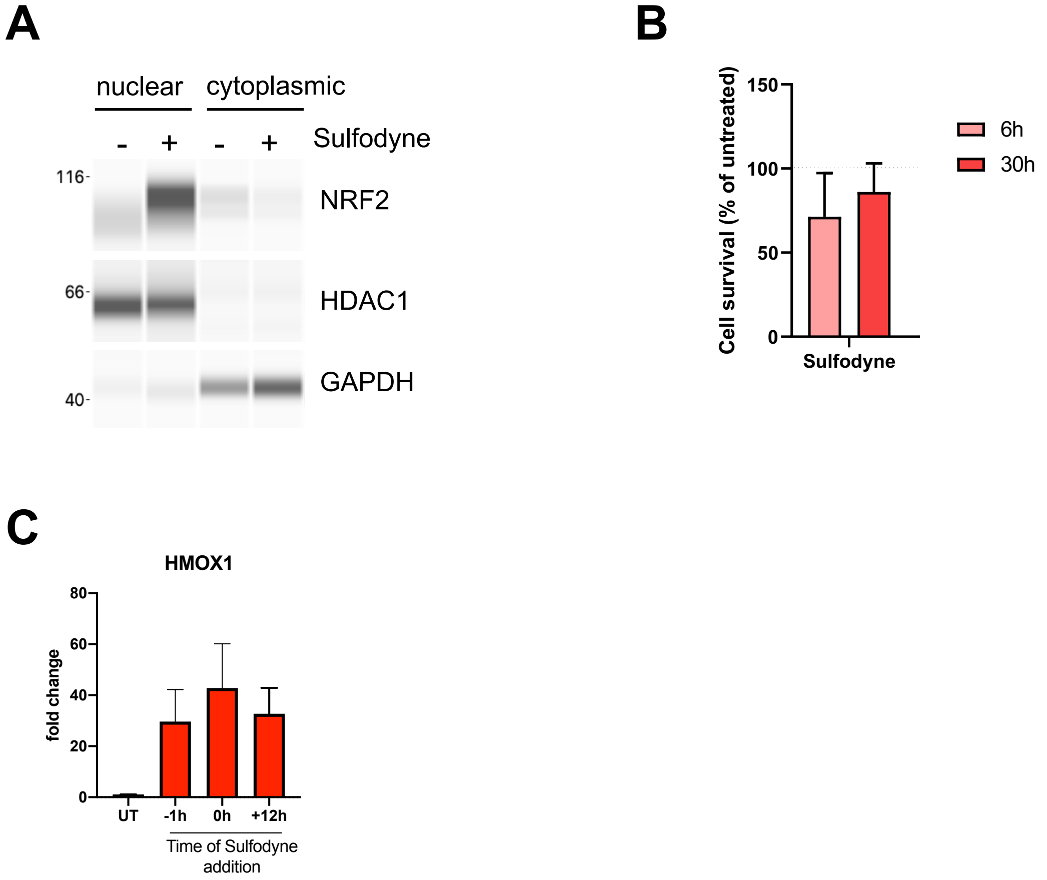

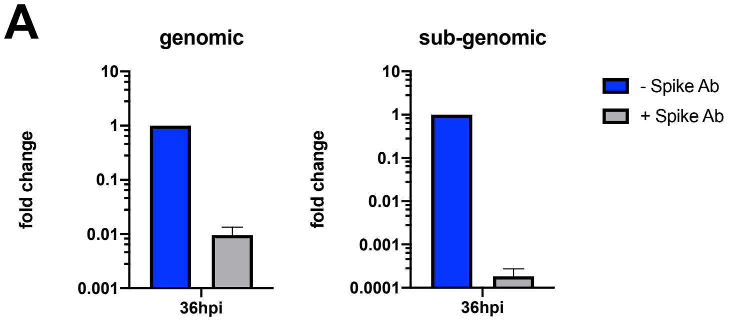

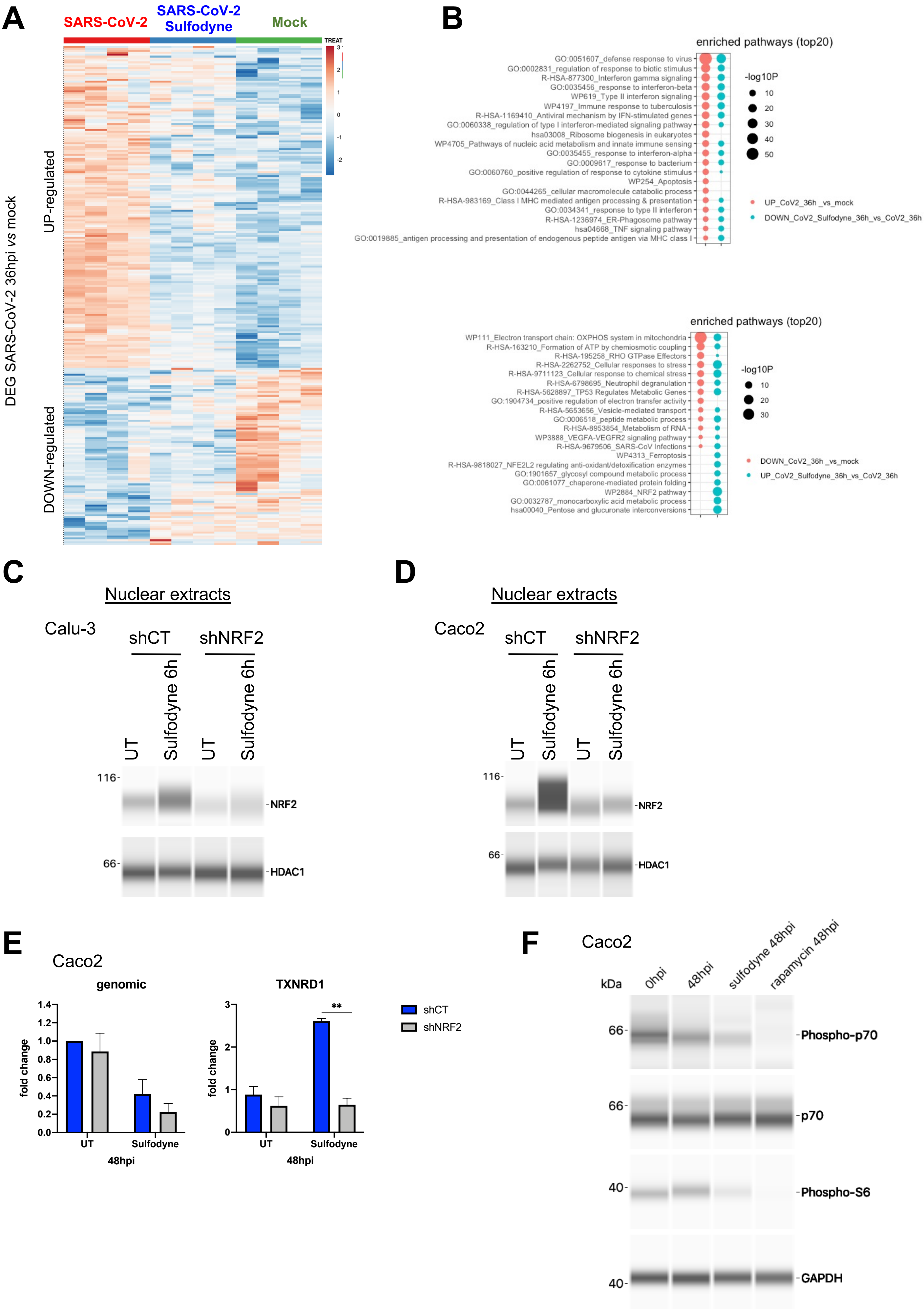

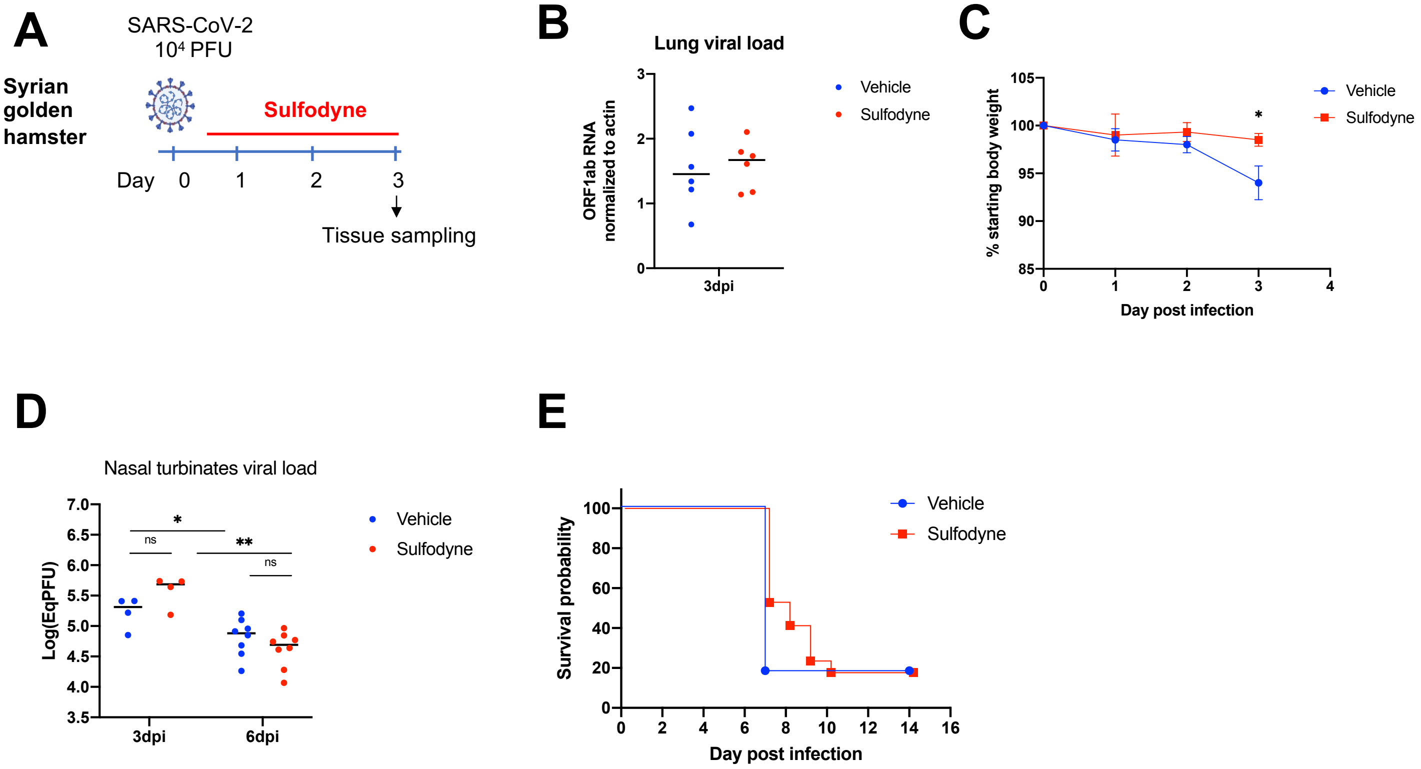
